# A parietal grid-like code rotates with cognitive maps but lags rapid behavioral transfer

**DOI:** 10.1101/2025.10.06.680746

**Authors:** Linda Q. Yu, Avinash Rao Vaidya, Aigerim (Aya) Akhmetzhanova, Sienna Bruinsma, Matthew R. Nassar

## Abstract

The neural grid code has been proposed to provide a mechanism for generalization and transfer of relational knowledge between situations enabling rapid adaptation of behavior in novel circumstances. However, to date, very little is known about the dynamics with which grid representations change at context transitions, or how such dynamics relate to downstream behavioral adaptation. Here we tested whether grid representations measured with fMRI rotate to match behavioral goals at context transitions and whether such rotations underlie knowledge transfer. Human participants performed a task that included unsignaled state changes at which the position of multiple target locations abruptly and synchronously rotated by the same degree. After state changes, participants were able to leverage the relative positions of the targets to rapidly infer locations, even novel ones, constituting a form of zero-shot transfer. We observed a cognitive grid-like code in the right posterior parietal cortex with a consistent phase angle that rotated with the relative positions of the targets. However, this rotation was too slow to account for rapid improvements in performance after a state change, and instead these improvements were more closely related to representations of the identity and location of spatial targets in the frontoparietal and orbitofrontal cortex, respectively. Our results highlight the ability of humans to rapidly transfer knowledge and demonstrate that a parietal grid-like code rotates into behaviorally relevant reference frames, but raise questions about the function of such rotations, pointing instead to alternate neural mechanisms for rapid knowledge transfer.

## Introduction

Humans and other animals are capable of generalizing from past experiences to novel situations they have never experienced before. Each time you visit a new grocery store, you are able to leverage past experience of having navigated similar environments to infer where certain items are likely to be relative to each other (e.g. fruits and vegetables are often adjacent). This capacity improves behavioral efficiency and reduces the need for tedious error-driven learning in each novel environment. Theoretical accounts have argued that the brain achieves this kind of behavioral transfer by maintaining a cognitive map of the environment that can be used for making inferences about such relationships (Eichenbaum & Cohen, 2014; Epstein et al., 2017; Tolman, 1948; Wilson et al., 2014).

Grid cells, first observed in the rat entorhinal cortex, are widely thought to support such an internal cognitive map. In rodents and in humans, these cells have been observed to fire in a triangular lattice pattern that tiles the spatial environment (Hafting et al., 2005; Jacobs et al., 2013). Recent studies in rodents and non-human primates have observed firing in entorhinal cortex (ERC) neurons that tiles more abstract task spaces defined by visual landmarks or the frequency of auditory cues (Aronov et al., 2017; Neupane et al., 2024). These findings are corroborated by human fMRI studies that have observed modulation in the BOLD signal in entorhinal cortex that follows an expected pattern based on a grid-like code when navigating virtual reality environments (Bellmund et al., 2016; Doeller et al., 2010; Kunz et al., 2015; Raithel et al., 2023), egocentric visual space (Julian et al., 2018; Nau et al., 2018), conceptual spaces (Park et al., 2021; Vigano et al., 2021), and other sensory perceptual spaces (Bao et al., 2019; Constantinescu et al., 2016). These studies have also identified signals consistent with a grid-like code outside of the entorhinal cortex, including in the piriform cortex, medial prefrontal cortex (mPFC) and posterior parietal cortex (PPC) (Bao et al., 2019; Constantinescu et al., 2016; Doeller et al., 2010; Raithel et al., 2023), as have direct electrophysiological recordings in humans (Jacobs et al., 2013). Together, this work has inspired a new perspective suggesting that the grid code may support navigation of internal conceptual spaces beyond spatial navigation (Behrens et al., 2018; Bellmund et al., 2018; Peer et al., 2021).

Previous work has observed that firing fields of grid cells in the entorhinal cortex rotate in concert when the animal is placed in a novel environment (Fyhn et al., 2007). This behavior is different from hippocampal place cells, which do not show any structured relationship between their firing patterns in different contexts (Muller & Kubie, 1987; O’Keefe & Conway, 1978). Recent work has suggested that changes to grid population codes can be rapid, occurring even after a single observation (Wen et al., 2024). These findings have inspired models and theoretical accounts that propose how grid codes capturing abstract structural regularities of the environment could be used to make predictions that guide behavior during context changes or in novel settings (Dong & Fiete, 2024; Whittington et al., 2020; Yu et al., 2021). For example, rotating a grid code onto which associations have been formed at one grocery store might enable automatic transfer of those associations to a new one. However, it remains unclear whether and how grid codes participate in supporting this kind of behavior.

Here we developed a novel paradigm to measure transfer learning in order to better understand the neural mechanisms that underlie it. In this paradigm, navigation target locations rotated together in concert, allowing us to specifically examine the rotational dynamics of the neural grid code over the course of trials, and to test its relation to concomitant transfer learning. While we observed evidence for a parietal grid-like code that rotated into a behaviorally relevant reference frame, we found that this rotation was too slow to facilitate the rapid behavioral improvement that occurs after a state change and was not predictive of transfer performance. Instead, we found representations of other task features in parietal and orbitofrontal cortex that more closely tracked behavioral performance including individual differences in transfer to novel contexts. Our results shed light on neural mechanisms of transfer learning and raise questions about the role of slower dynamics of neural grid codes.

## Results

### A predictive inference task to measure behavioral transfer

Across three experimental sessions, participants completed several variants of a predictive inference task that was framed in terms of blocking alien missile attacks. On each trial, participants were asked to move a “shield” (colored circle) to block an incoming “alien attack” (red dot) that would occur at some location on a large circle depicting a “planet” (Figure 1a). Attack locations were drawn from a 2D gaussian distribution centered on target locations that were specific to each shield color. Trials were organized into “mini-blocks” containing each shield color presented in random order. Target locations for each shield color remained fixed for 6-8 mini-blocks (a “state-block”), after which all target locations rotated conjointly without any warning (“state changes”; Figure 1b). In sessions one and two, such rotations alternated between +90 and -90 degrees giving rise to two distinct target location configurations. In session three rotations could occur to any of 23 possible target configurations that resulted from rotating a single set of target locations in 15 degree increments up to 345 degrees (Figure 1c). Sessions two and three included behavioral tests of generalization: in session two this test came in the form of novel colors (i.e. can participants put a new shield in the correct location after only observing it in the alternate rotation?) whereas in session three this came through the novel rotations (i.e. after participants see that the target location for one of the colors has rotated to a completely new position, can they place the other colored shields appropriately?) Sessions one and three included only behavior whereas session two was performed in an MRI scanner while functional neuroimaging data were collected.

**Figure 1.**
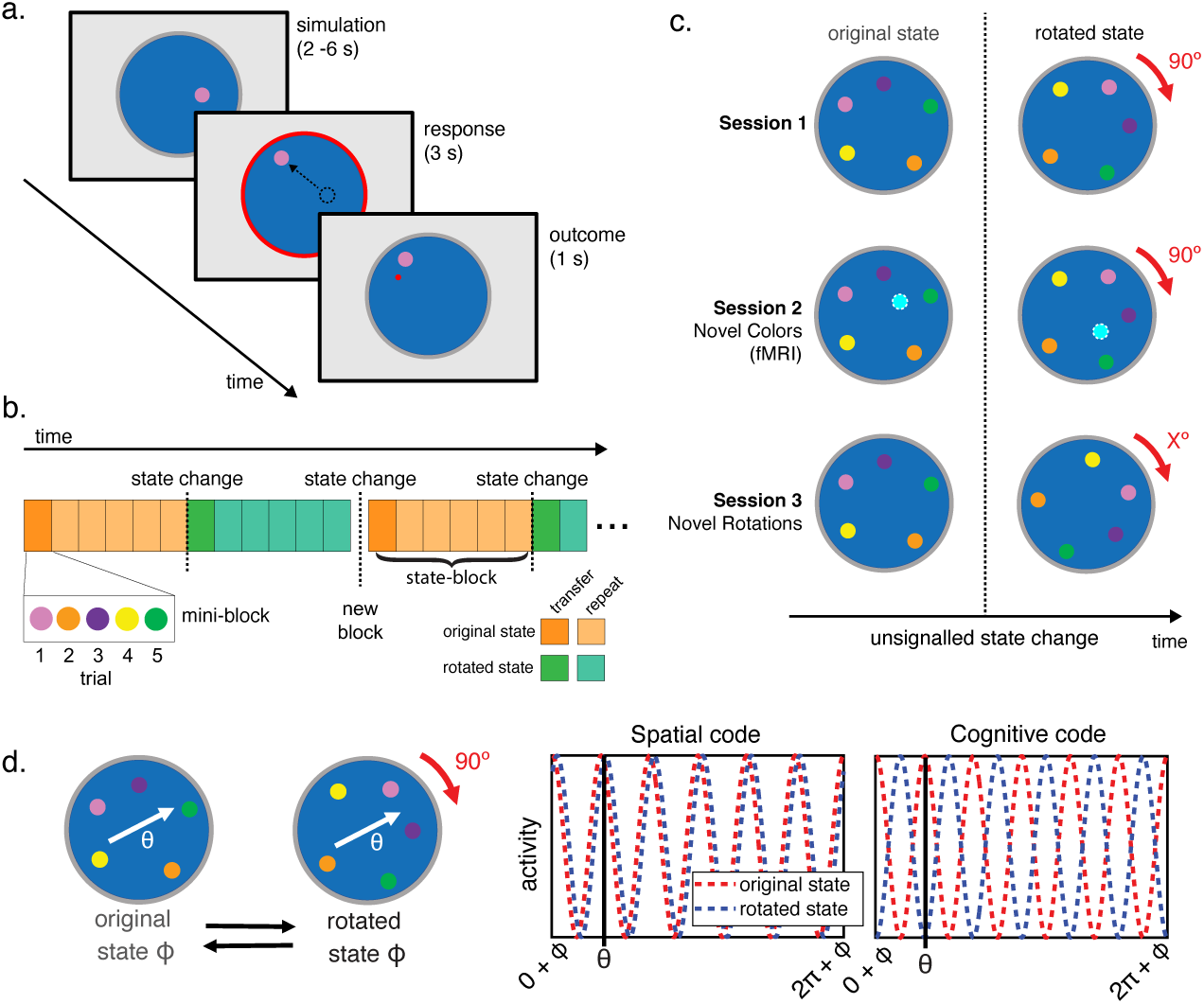
Measuring transfer learning with a predictive inference task. **a.** Participants moved a “shield” (colored circle) to a location where they believed it would block an upcoming “alien attack” (red dot). Alien attacks were normally distributed around a target location that di>ered for each shield color. On each trial, a colored shield would appear in a random position on the screen and participants were asked to mentally simulate their planned movement. Participants were then cued to start moving the shield when the border of the planet turned red. The location of the alien attack was then shown at the outcome phase of the trial, either as a red dot if the participant missed the attack or as a green dot if the participant successfully blocked the attack. **b.** Schematic of task timeline. Participants completed trials with all shield colors in a randomized order (except for novel colors in session 2). After several such mini-blocks within the original state, the target locations all rotated in concert at a state change. At the start of each task block, the state would change again, either back to the original state (in sessions 1 and 2), or to a new rotation state (session 3). **c.** Each session included unsignaled state changes at which all target locations would all rotate in concert with respect to the center of the planet. In session 1 and 2, all target locations would rotate 90 degrees clockwise to the rotated state, and then rotate 90 degrees counterclockwise back to the original state on the subsequent block. In session 2, participants were also presented with novel color shields, each presented only in two state-blocks of the task, one for each rotation condition, and each with their own unique target location. This condition required participants to transfer the learned 90 degree rotation rule to new cases. In session 3, novel rotations were introduced where the target locations for each shield color rotated by di>erent amounts from the original position at each state change, requiring participants to apply new rotation rules to the previously learned target locations. **d.** Schematic illustrating cross-validation test for identifying a cognitive or spatial grid like code in a region of interest (ROI). Two grid angles (ɸ) are separately estimated using regressors for the six-fold sine and cosine of the traversed angle (θ) in both the original state and the 90° rotated state. Each grid angle is then used to create a regressor for the predicted activity in the other state, based on the trajectories in each trial. For a region with a grid-like code that is fixed to the real space of the display (i.e. a ‘spatial’ code), the grid angle for the two states should be similar and the predicted activity for any traversed angle (θ) should be positively correlated in the two states. However, if the grid angle is fixed to the relative positions of the colors (i.e. a ‘cognitive’ code), then the predicted activity for a given trajectory in the two states should be negatively correlated.

This study was designed to specifically test whether rotation of the grid code during these state changes supports transfer behavior. In order for a grid representation to transfer stored behavioral policies from one rotation condition to another, the entire grid code would need to rotate (reorient) across these conditions, such that the representation of a given color target location on the grid code remains constant across all conditions. We were thus interested in whether any region displayed properties of this sort of ‘cognitive’ grid code that is fixed to the relative target positions for each shield color rather than the physical space of the display. To test this, we carried out a between-state cross validation test, adapting the method developed by Doeller et al. (2010) (Figure 1d). We fit a grid angle in each rotation state separately using six-fold sine and cosine regressors on the traversed angle. We then created regressors centered on these grid angles to test in the held-out state. In this analysis, if a brain region carries a *spatial* grid-like code fixed to the physical display, then this regressor should be positively correlated with the BOLD signal in the held-out state as the grid angles fit in the two states should be in-phase. Indeed, this is typically the prediction for a region reflecting a consistent grid-like spatial code (Bao et al., 2019; Constantinescu et al., 2016; Doeller et al., 2010; Park et al., 2021; Vigano et al., 2021). However, if a region carries a *cognitive* grid-like code fixed to the relative mean target positions for each shield color then, because the rotated state is 90° from the original state, the grid angles for the two states should be precisely antiphase (i.e. oriented 30° apart), and the regressor should be negatively correlated with the BOLD signal in the held-out data. Thus, our task and analysis allowed us to identify either visuo-spatial or cognitive grid representations that maintained a consistent phase across rotations.

### Participants learned relational structure and used it to make inferences

Participants rapidly learned to place shields in appropriate locations during their first behavioral session, benefitting from experience over multiple timescales. We examined how participants’ performance changed across mini-blocks, blocks, and the entire task using a normalized error term that quantified actual errors (distance between final shield position and target location) relative to errors computed according to the target location that would be appropriate for the alternate rotation state (distance between final shield position and target location in the alternate rotation; Figure 1c). Participants displayed rapid behavioral adjustments within the first mini-block after a state change, before any of the colors had been repeated, consistent with transferring knowledge about a rotation observed for one color to update predictions for all others. Participants’ normalized error dropped rapidly over these first few trials after a state transition (main effect of trial number: t(47) = 11.15, P < 0.0001, d = 1.61) and this effect was most pronounced later in the session, once participants had experienced several rotations (interaction of trial and block number: t(47) = 4.80, P < 0.0001, d = 0.69; Figure 2a).

**Figure 2.**
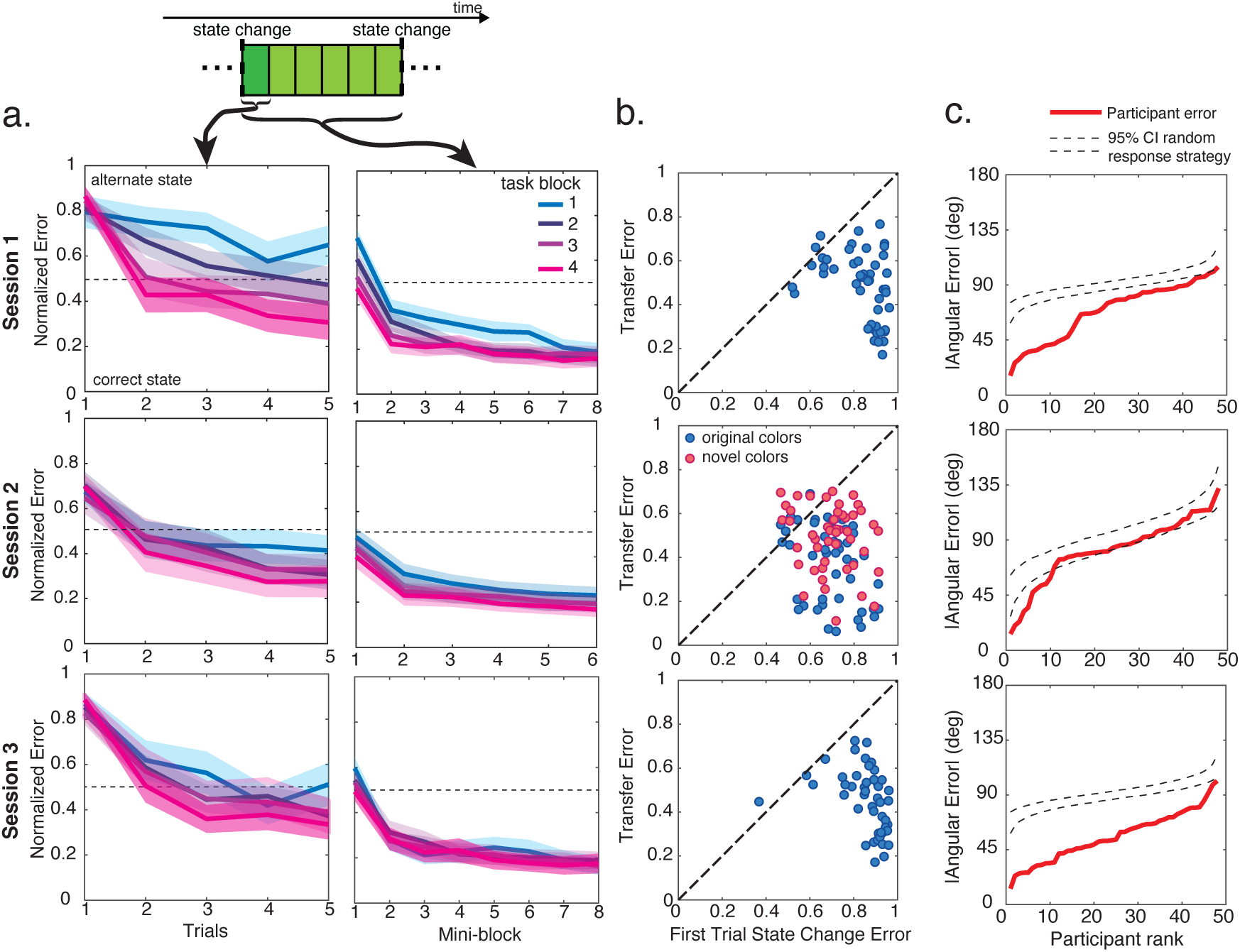
Behavioral measures of learning and transfer. **a.** Left: Normalized error on transfer trials in sessions 1-3 (the first mini-block after a state change) is plotted for the first 5 trials after a rotation. Note that each of these trials reflects the first time seeing the new shield color after the rotation. Right: normalized error as a function of mini-blocks after each state change is shown in the right panels. Solid colored lines represent means in each block, shaded area represents the 95% confidence interval. **b.** Scatter plots for individual participants showing mean normalized error during sessions 1-3 for the first trial after the state change against normalized error on first presentation for other colors (i.e. transfer trials). **c.** Absolute angular error for transfer trials in sessions 1-3 ranked by participant error. Dashed lines show the 95% CI for the expected error following a random response strategy in simulated data. Session 2 panel shows data from novel color trials only.

Participants also showed performance improvements at a longer time scale, measured as reduced normalized errors, across task blocks (main effect of block: t(47) = 7.18, P < 0.0001, d = 1.04) and across mini-blocks after each state change (main effect of mini-block: t(47) = 19.91, P < 0.0001, d = 2.87). The improvement across mini-blocks became less pronounced over blocks as participants reached asymptotic performance faster (interaction of block and mini-block: t(47) = 6.86, P < 0.0001, d = 0.99). Thus, performance in the first task session demonstrated learning across multiple timescales and suggested transfer of rotational knowledge across different stimulus categories.

To more directly test if participants capitalized on relational structure to infer color target locations, we focused on trials 2-5 in the first mini-block after a state change. These trials, which we refer to as “transfer trials” reflected the first time a participant saw a given shield color in the state-block and could conceivably infer that a rotation had occurred (based on the feedback from the first trial after the rotation). Participants’ normalized error on these trials was significantly below their error on the first trial after a state change (t(47) = 10.46, P < 0.0001, d = 1.51; Figure 2b). Participants’ error was significantly above the expected error given a random response strategy in the first block of the task (normalized error of 0.5; two-tailed one sample t-test: t(47) = 2.71, P = 0.009, d = 0.39), but dropped significantly below this level by the last block (t(47) = 2.60, P = 0.01, d = 0.38). Thus, while participants persisted in the original state after a state change at the start of session 1, by the end of the session they were able to adapt to these state changes after just a single trial.

To characterize how many participants were able to adaptively move the shield to its alternate state location in these transfer trials, we compared each participant’s mean absolute angular error on transfer trials to the expected angular error for a random response strategy (Figure 2c). Participants’ angular error on transfer trials was well below the 95% CI of a simulated null distribution for all but one participant, indicating the vast majority showed behavioral evidence of successful transfer.

Task performance in the second session further supported that participants used relational structure to make inferences. Like in session 1, we observed that participants’ normalized error for the original colors dropped over trials within the first mini-block (t(47) = 11.45, P < 0.0001, d = 1.65), and over mini-blocks after each state change (t(47) = 14.08, P <0.0001, d = 2.03). Participants’ normalized error on transfer trials was significantly below that of the first trial after the block transition (two-tailed, within-subjects t-test: t(47) = 9.08, P < 0.0001, d = 1.32), and significantly below the error expected for random responding (i.e. normalized error of 0.5: t(47) = 4.77, P < 0.0001, d = 0.69) (Figure 2b). Participants’ absolute angular error on original color transfer was below the 95% CI for error expected given a random response strategy in all participants, demonstrating successful transfer behavior.

We next carried out the same tests for the novel colors. Notably, participants had only practiced moving the shield to the target position in one state for these novel colors, and thus were forced to use their knowledge of the target position in that trained state along with its relationship to the other target locations in that state to infer the correct position in the new state. Normalized error on novel color transfer trials was reduced relative to the first trial after a state change (t(47) = 6.95, P < 0.0001, d = 1.00; Figure 2b). However, normalized error was not reliably below chance on the normalized state error metric (t(47) = 0.41, P = 0.68), raising questions as to whether participants had improved their error metric by simply guessing on the novel color rotation trials. To disambiguate inference versus random guessing at the level of individual participants, we constructed the null distribution of rank ordered mean absolute errors expected if all participants were guessing and compared our data to this null distribution. We found that the best performing 10 participants (20.8%) had performance incompatible guessing on the novel color trials, whereas the remaining participants fell within the 95% CI of the null distribution ( Figure 2c). Thus, while many participants resorted to guessing for the completely new colors, a minority demonstrated clear evidence of successful transfer in these trials. Consistent with this idea, examining the distribution of errors across participants revealed a mostly uniform guessing strategy in poor performers, and a mixture of guessing and correct target inference for these novel colors in the top performers (Supplementary Figure 1).

Behavioral data from the third session demonstrated that all participants were able to transfer relational information about the shield colors to novel rotations (Figure 2a). Unlike in the first two sessions in which participants were extensively trained on two rotation conditions, each rotation in the third session was completely unique. Thus, performing well on transfer trials in the third session required participants to place the shield for a given color based only on feedback observed for one or more other shield colors. Performance after novel rotations was, as expected, impaired for the first trial following a novel rotation (two-tailed t-test against normalized error of 0.5: t(46) = 21.78, P < 0.0001, d = 3.21). However, performance improved markedly for subsequent transfer trials, even though the shield colors from these trials had not yet been associated with their current target location (two-tailed t-test against normalized error of 0.5: t(46) = 2.29, P = 0.03, d = 0.34; Figure 2b). The absolute angular error on transfer trials was below the 95% CI for the expected error given a random response strategy for all participants (Figure 2c).

These results demonstrate that participants were able to transfer knowledge about the relational structure across target locations to correctly place colored shields in completely new locations. Thus, a primary question arising from the observed behavioral data is how the brain accomplishes this sort of flexible transfer.

### Right posterior parietal cortex maintains a cognitive grid-like code

To better understand the neural mechanisms through which people generalized knowledge across rotations, we examined fMRI data collected in session two, and in particular tested the degree to which behavioral transfer might have been afforded through rotation of grid-like codes (Whittington et al., 2020; Yu et al., 2021). The regular triangular lattice structure of the grid code is thought to yield a hexagonally symmetric change in the BOLD signal with respect to participants’ angle of movement (Doeller et al., 2010). We thus first sought to identify regions of the brain where the BOLD signal exhibited a hexagonal symmetry with respect to the angle along which participants moved the shield in each trial. We adopted an approach used in previous studies to localize such signals by carrying out an F-test on sine and cosine regressors with 60° periodicities (Bao et al., 2019; Constantinescu et al., 2016; Park et al., 2021). To adapt this method to the current paradigm, we included two pairs of these regressors for the two different task states and carried out this test on both pairs. This analysis identified a set of brain regions that have been found to exhibit such a hexagonally symmetric signal in other fMRI studies (Bao et al., 2019; Constantinescu et al., 2016; Doeller et al., 2010; Park et al., 2021), including the mPFC, posterior cingulate cortex (PCC), left and right temporoparietal junction and right posterior parietal cortex (rPPC) (Figure 3a). Consistent with prior work (Constantinescu et al., 2016), we focused this analysis on the simulation period when participants are planning their trajectory rather than the period of actual movement to avoid potential motor confounds. However, similar results were observed when we carried out the same analysis on the response period (Supplementary Figure 2).

**Figure 3.**
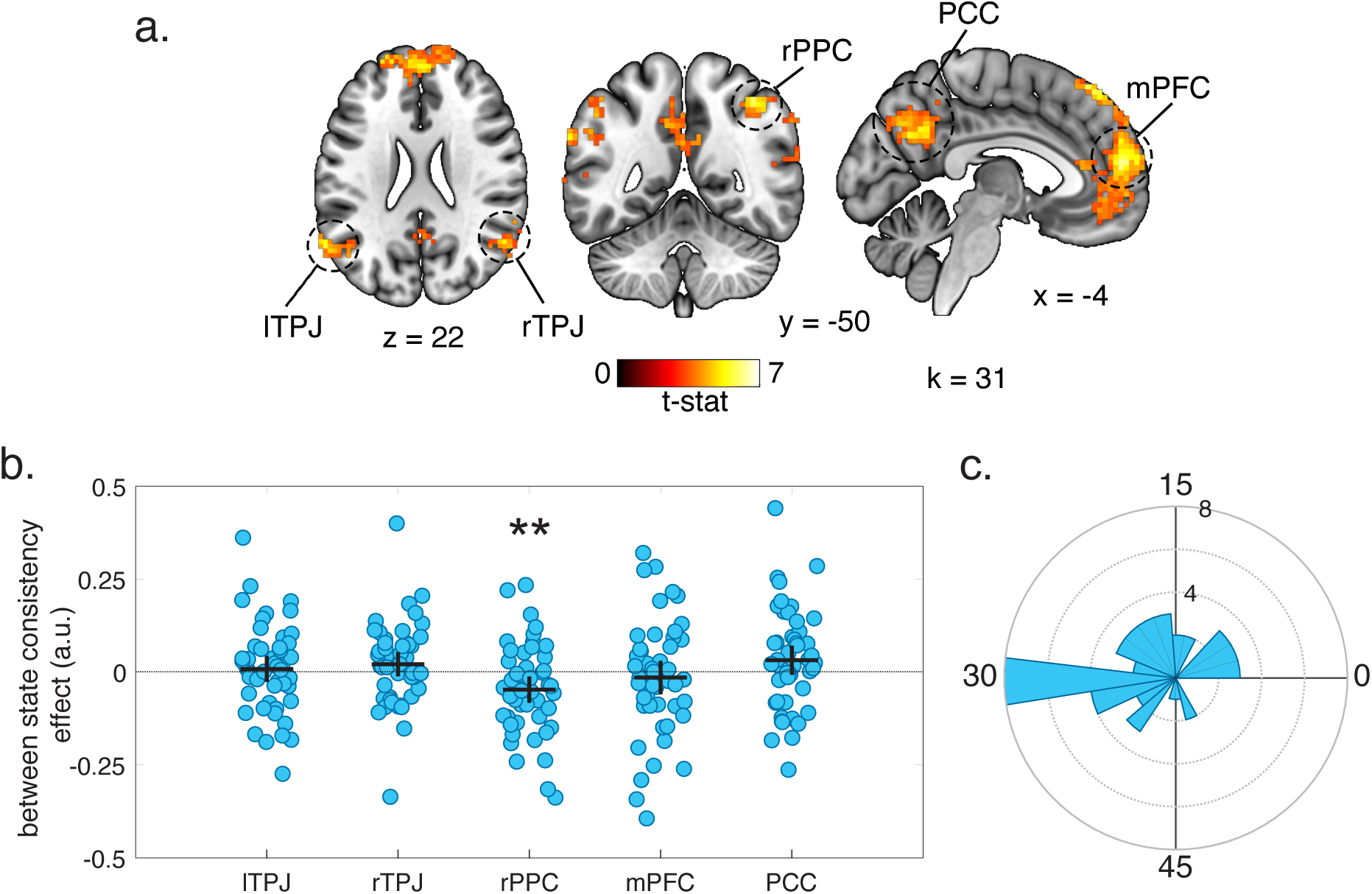
Identifying brain regions with a grid-like cognitive or spatial code. **a.** Statistical map from whole-brain analysis identifying regions where activity is modulated by a hexagonally symmetric signal in both the rotated and original states. This analysis identified such a signal in the left and right temporal parietal junction (TPJ), right posterior parietal cortex (rPPC), posterior cingulate cortex (PCC) and medial PFC (mPFC). Results are displayed at a cluster forming threshold of P<0.001 and corrected for multiple comparisons with permutation tests for defining a cluster extent threshold (k) at P<0.05. **b.** CoeWicients for between-state consistency test for ROIs identified by whole-brain test in a. Markers represents individual participants and Horizontal/vertical bars represent mean/95* CI. Negative between-state coeWicients in the rPPC indicate that this ROI maintains a cognitive grid-like code (two-tailed one-sample t-test: t(47) = 2.72, P = 0.009, d = 0.39). **c.** Polar histogram for the angular diWerence of grid angles estimated in the original and rotated state in the rPPC across participants.

A cross-validation test on the grid angles for ROIs identified in the whole brain F-test revealed evidence for a cognitive grid-like code in the rPPC. As described above, we carried out a cross-validation test by separately fitting a grid angle in the two different rotation states and testing this angle in the other (Figure 1d). We carried out this test on spherical ROIs defined around the peak coordinates of those regions identified as exhibiting a hexagonally symmetrical signal in the whole-brain F-test. Coefficients in the rPPC were significantly below zero (two-tailed, one-sample t-test: t(47) = 2.72, P = 0.009, d = 0.39), consistent with predictions for a cognitive grid-like code (Figure 3b). We compared the difference between grid angles in the two states to confirm that most of angles were oriented approximately 30° apart (Figure 3c). The grid angles for the two states were more than 15° apart in the majority of participants (73%). No other ROI tested showed evidence for maintaining either a consistent spatial or cognitive grid-like code (Figure 3b).

Follow up validation tests suggested that the cognitive grid code in the rPPC was specific to a six-fold symmetry characteristic of hexagonal grid coding in the brain (Figure 4a). We did not observe evidence for grid-like code with any other fold symmetries in between-state cross-validation tests aside from the canonical six-fold symmetry expected for grid (Figure 4b; Doeller et al. (2010).

**Figure 4.**
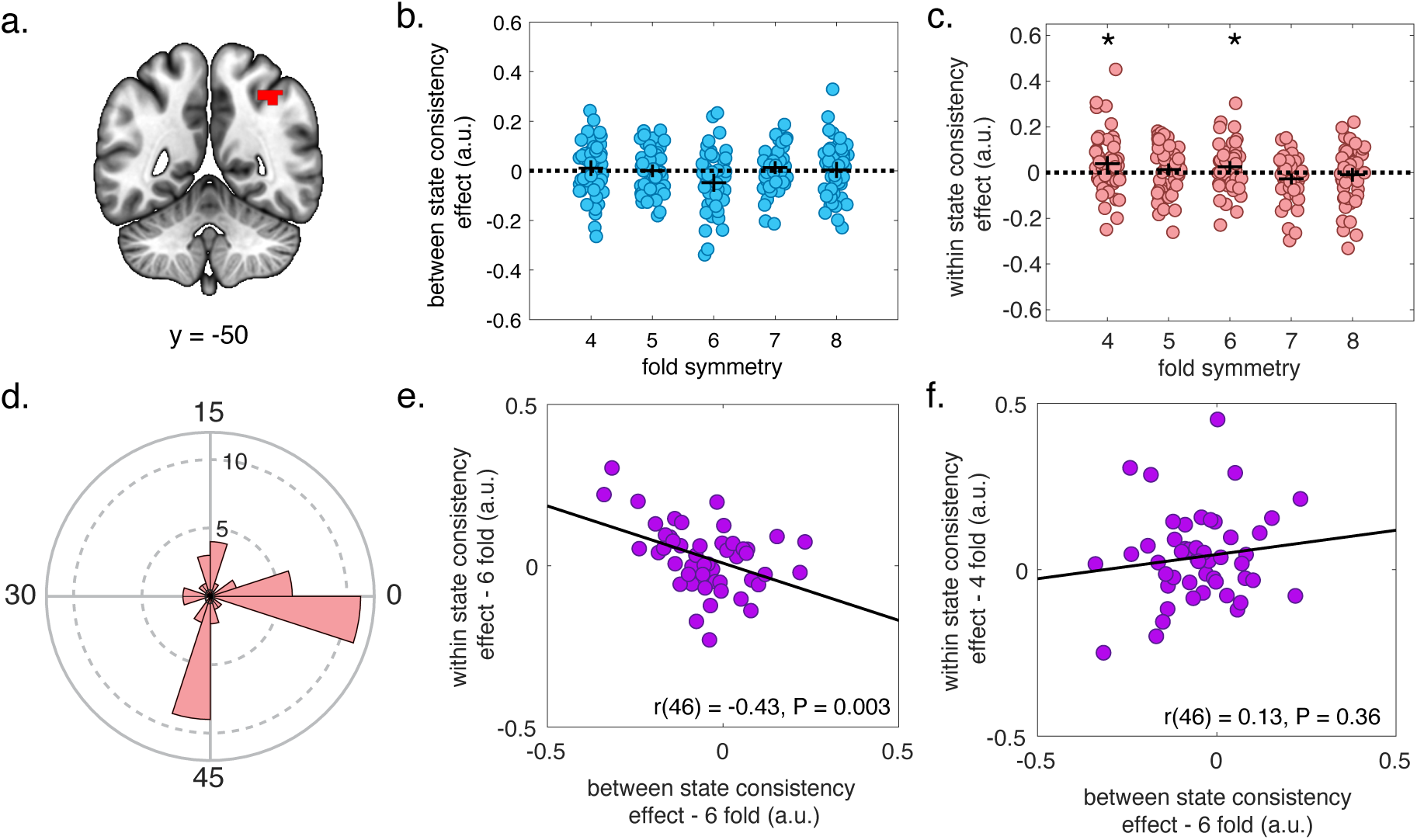
Validation tests for right posterior parietal cortex (rPPC) grid-like code. **a.** rPPC ROI. **b.** Participant coefficients from between-state consistency test for predicted signals with different fold symmetries showed no other significant effects aside from the 6-fold code (same data shown in Figure 2c) **c.** There were significant effects for both the 4-fold and 6-fold regressors when testing for the consistency of this grid-like code within the same states. Horizontal bars represent the mean, vertical bars represent the 95% CI. *P < 0.05, right-tailed, one-sample t-test against zero. **d.** Polar histogram showing the distribution across participants for the mean difference between grid angles from each run against the mean grid angle estimated from all other runs within each state. **e.** There was a significant negative relationship between rPPC coefficients from between-and within-state consistency tests across participants for the 6-fold regressor, consistent with participants with a stronger cognitive grid-like code in this region also showing evidence for more consistent code within the same state. **f.** There was no equivalent significant correlation between the within-state coefficients for the 4-fold grid code and the between-state coefficients for the 6-fold regressor in the rPPC.

The rPPC grid-like code was internally reliable – and its reliability across participants was related to the strength of the cognitive rotation between states. We anticipated that if the rPPC maintains a stable cognitive grid-like code with antiphase grid angles between the two states, then grid angles should be in-phase for repetitions in the same state. To test this, we fit grid angles for both states within each scanner run (one task block) separately and used a leave-one-out cross-validation procedure to test whether a regressor centered on the mean grid angle across a set of runs could independently predict activity in this ROI in the held-out run. Coefficients from these regressors were significantly greater than zero (right-tailed, one-sample t-test: t(47) = 1.71, P = 0.047, d = 0.25) for the six-fold regressor (Figure 4c). The grid angles from these different runs were less than 15° apart in a slight majority of participants (58%; Figure 4d). Unlike the results from our rotational phase-locking analysis (Figure 4b), the internal reliability of rPPC was not specific to 6-fold symmetry. We also observed a significant effect for a regressor constructed with four-fold symmetry (Figure 4c; right-tailed, one-sample t-test: t(47) = 2.02, P = 0.025, d = 0.29).

Nonetheless, further investigation supported that the cognitive grid representations observed in rPPC were more tightly related to six-fold grid like codes, rather than any stable four-fold spatial encoding. We anticipated that if the rPPC maintains a cognitive six-fold grid code, then the strength of the within-state consistency effect for the six-fold regressor should be inversely correlated with the between-state effect (where a more negative effect indicates more consistent antiphase coding). We found such an inverse relationship for the coefficients from the six-fold within-state cross-validation test (Figure 4e; r(46) = -0.43, P = 0.003), but not for coefficients from the four-fold cross validation test (r(46) = 0.13, P = 0.36), arguing for the validity of the observed grid-like code in the rPPC (Figure 4f).

We did not observe a reliable grid-like code in the ERC. While our whole-brain analysis identified a similar network of regions displaying hexagonal modulation as previous studies (Bao et al., 2019; Constantinescu et al., 2016; Doeller et al., 2010), one notable distinction between our study and previous fMRI grid representation findings is that we did not find any evidence of a hexagonally symmetric signal in the ERC even at a very liberal threshold of P < 0.05, uncorrected (Supplementary Figure 3). We also did not find evidence for a consistent grid-like code in the ERC cross-validation tests (Supplementary Figure 4), though evidence for such a grid-like code in the ERC was positively correlated with that in other ROIs identified in this whole brain analysis (Supplementary Figure 5).

### rPPC cognitive grid-like codes rotate too slowly to account for rapid behavioral transfer

Given that grid-like codes in rPPC were specific, reliable, and formatted in a way that could potentially transfer knowledge across conditions, we next sought to test whether they could explain inter-individual and trial-to-trial differences in transfer behavior.

We found no evidence that the cognitive grid-like code in rPPC was related to individual differences in task performance. We computed correlations between the between-state consistency of the six-fold grid-like code in rPPC and normalized error on transfer trials with original colors, novel colors, and on novel rotations. In all cases, we found no evidence of a significant relationship (all P’s ≥ 0.2; Supplementary Figure 6).

Indeed, the direction of the effect was the opposite to the one consistent with cognitive grid codes mediating improved behavior in all cases. Similarly, we found no relationship across participants between performance on transfer trials and the mean of between-state coefficients in the bilateral ERC and other ROIs identified in the hexagonal symmetry analysis (all P’s ≥ 0.6).

We observed a relationship between the trial-wise strength of a cognitive grid-like code in rPPC and behavior, but this was driven largely by the fact that cognitive grid-like code changed as a function of trials within a state. We adapted a method used by Julian and Doeller (2021) to derive estimates of which state (i.e. the rotated or original state) the signal in this region was consistent with at a trial-wise level. Using the grid angles from the between-state cross-validation analysis, we generated regressors for the expected response in each trial for the alternate state and estimated Pearson correlation coefficients between these regressors and the BOLD signal in this ROI, after removing linear effects for trial and movement regressors. This provided us with a trial-wise measure of the strength of evidence for a grid code that was either in-phase or antiphase with the alternative state. We found a small, significant relationship between these coefficients and normalized error across trials in a regression analysis on all trials, where trials with more positive coefficients were associated with greater normalized error (one-tailed, one sample t-test: t(47) = 1.79, P = 0.04, d = 0.26). This modest effect might be interpreted as evidence for a relationship between grid alignment and behavior, a necessary condition for the idea that grid rotations underlie the knowledge transfer in our task. However, when we included mini-block number as a covariate in the regression model, the effect was eliminated (t(47) = 1.18, P = 0.2, d = 0.17), suggesting that the relationship between error and single-trial coefficients was the result of a common relationship with the number of elapsed mini-blocks.

Examining the dynamics of rPPC grid representations revealed that they rotate far too slowly to account for rapid and flexible behavioral transfer. We tested the relationship between these trial-wise coefficients and trial number on transfer trials after a state change (i.e. on the first mini-block), and across mini-blocks. There was no change in the trial-wise correlation coefficients over trials after a state change on the first mini-block (Figure 5a; two-tailed, one-sample t-test: t(47) = 0.42, P = 0.67, d = 0.06). However, these correlation coefficients did become significantly more negative over the course of mini-blocks following a state change (Figure 5b; two-tailed, one-sample t-test: t(47) = 2.16, P = 0.036, d = 0.31). This drop was only evident by the third mini-block, corresponding to trials 15-21 after a rotation, by which time participants’ mean normalized error had already reached asymptotic levels. This cognitive grid-like code thus appeared to evolve more slowly than would be required for a neural signal underlying the rapid behavioral improvement observed on transfer trials in this task.

**Figure 5.**
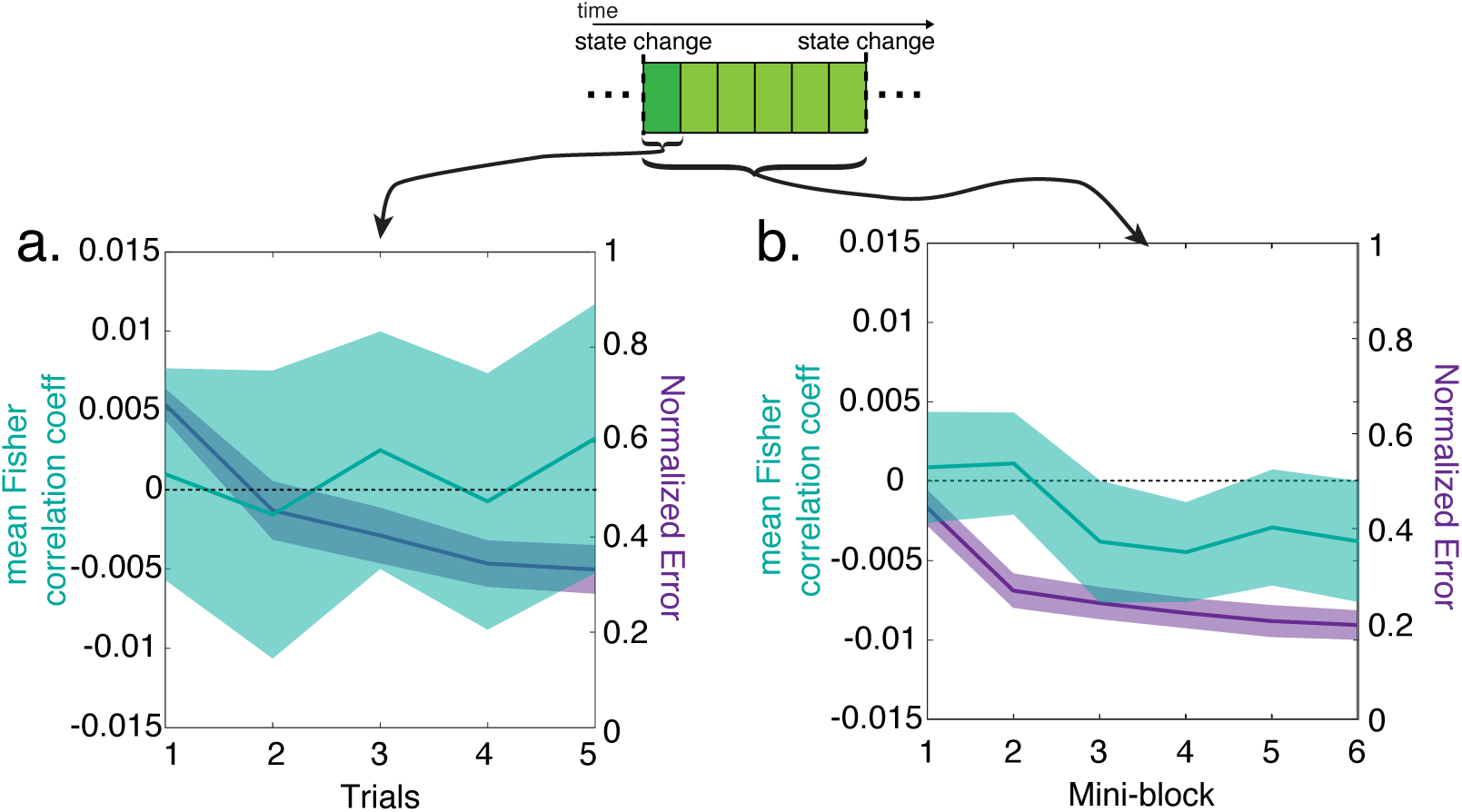
Trial-wise analysis of grid-like code phase consistency between states in the right posterior parietal cortex (rPPC). Mean Fisher transformed correlation coefficients reflect the degree to which the BOLD signal correlated with single trial regressors based on the grid angle fit in the alternate state. Here, positive correlation coefficients indicate a reliable spatial grid code shared between the two states, while negative correlation coefficients indicate a reliable inverse relationship between the expected activity in the two states, consistent with a cognitive grid code. **a.** Mean Fisher transformed correlation coefficients from trial-wise phase consistency analysis and normalized error as a function trial during the first mini-block after a state transition **b.** The same as **a**, as a function of mini-block. Note that behavioral evidence of transfer precedes changes in the grid angle phase by several mini-blocks. Lines represent means with shaded area representing the 95% CI.

Comparing this trial-wise measure of the grid-like code in trials sorted based on participants’ behavior further argued against a relationship to behavior. We observed no significant differences in these single-trial coefficients between trials where participants moved the shield into the correct position for the current versus the alternative state (Supplementary Figure 7). Thus, while the rPPC grid code was specific, reliable, and formatted in a way that could potentially afford transfer, its timing was too slow to account for rapid improvements in behavior, suggesting that the brain achieved this fast transfer through other means.

### Representations of target identity and spatial position during simulation predict transfer behavior

We next tested for other potential alternative representations that might support transfer behavior. To do so, we used a trial-wise whole-brain representational similarity analysis (RSA) to identify correlates of task variables that could putatively support transfer behavior. We estimated a t-statistic map for the simulation phase in each trial (Mumford et al., 2012) , and calculated the correlational distance between voxels in each trial to make a trial-by-trial neural representational dissimilarity matrix (RDM). We use a searchlight approach, where we carried out a multiple linear regression to compare the neural RDM with multiple hypothesis RDMs of interest at each pass, including: 1) a shield identity RDM where trials of the same color shield were similar to each other and uniformly dissimilar from all other trials (Figure 6a), 2) an RDM for the Euclidean distances of the mean target positions of each shield color in each state (Figure 6b).We also included several covariate RDMs to control for factors that were not of interest (e.g. shield starting position, the estimated perceptual distance between colors, etc, see Supplementary Figures 8-10), but these were not significantly correlated with pattern activity in any regions, or were not related to transfer behavior.

**Figure 6.**
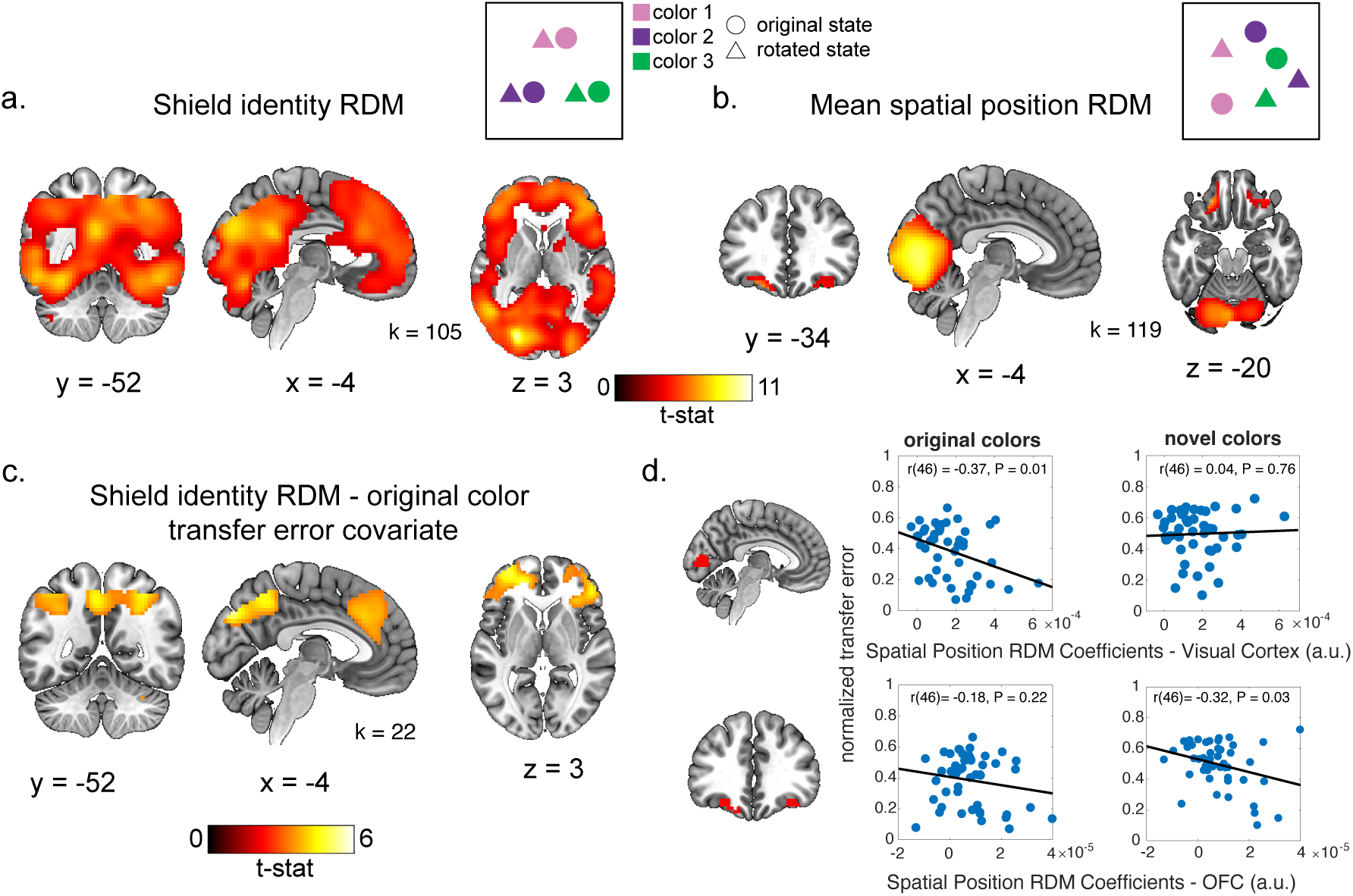
Representational similarity analysis and relationship of representations with behavior. **a.** Whole-brain group-level statistical map from searchlight analysis testing relationship of neural representational dissimilarity matrix (RDM) with a shield identity RDM, in which trials with the same shield color are similar to one another and uniformly dissimilar from all other trials (as illustrated in inset schematic at the top of panel). **b.** Group-level statistical map from whole-brain searchlight analysis for mean spatial position RDM, in which trial distances depend on the Euclidean distance between the mean positions of each shield color in the rotated and original state (see inset schematic). **c.** Statistical map for covariate analysis testing where there was a significant inverse relationship between the strength of the shield identity RDM representation and normalized error on transfer trials. This analysis was constrained to voxels where the shield identity RDM was significantly correlated with the neural RDM, corrected for multiple comparisons (shown in panel a). All statistical maps were defined with a cluster forming threshold of p<0.001 and corrected for multiple comparisons permutation tests to construct a null distribution over cluster extent. Cluster extent threshold corresponding to p=0.05 for each contrast is given by the value of k. **d.** Beta coefficients from the mean spatial position RDM in ROIs defined around peaks from whole-brain statistical map in panel b (abscissa) are plotted as a function of behavioral measures (ordinate) for individual subjects (points). The mean spatial position representation was significantly inversely correlated with transfer error for original colors (left top), but not novel colors (right top), while the opposite was true for an ROI defined in the bilateral orbitofrontal cortex (OFC; bottom).

Strength of the shield identity RDM in a frontoparietal network predicted transfer performance for original, but not novel colors. Several brain regions encoded shield identity consistently across rotation conditions including visual, parietal and prefrontal regions (Figure 6a). Given the large fraction of cortical regions encoding shield identify, we carried out an additional second-level test including behavioral variables as covariates of interest within a mask defined by voxels where we found an effect for the shield identity RDM at a cluster-corrected threshold. We found that error on transfer trials for original colors was inversely correlated with the strength of the shield identity RDM effect within the frontoparietal network, including rostrolateral prefrontal cortex, the bilateral PPC, posterior medial parietal cortex and dorsomedial prefrontal cortex (Figure 6c). However, there was no significant relationship between this shield identity RDM effect and error on novel color transfer trials that survived multiple comparisons correction. Thus, individuals who were able to successfully transfer knowledge to previously observed shield color better discriminated shield identity within a wide frontoparietal network.

The strength of the mean spatial position representation in the OFC and visual cortex also predicted transfer performance. The mean spatial position RDM was significantly correlated with the neural RDM in the visual cortex and bilateral orbitofrontal cortex (OFC) (Figure 6b). Across participants, this mean spatial position effect was inversely related to error on novel color trials in an ROI defined around the peaks of the OFC effects (r(46) = -0.32, P = 0.03, but not on transfer trials with the original colors (r(46) = -0.18, P = 0.2). This correlation between the spatial mean position effect and transfer performance on novel colors was not significant, but trending after controlling for the linear relationship between error on these novel color trials and repeat trials for the original colors (r(46) = -0.26, P = 0.069; Supplementary Figure 11a). The opposite pattern was found in the visual cortex, where the strength of the mean spatial position effect was inversely correlated with error on transfer trials with the original colors (r(46) = -0.37, P = 0.01), but not novel colors (r(46) = 0.04, P = 0.77). Thus, the strength of spatial target coding in OFC, but not visual cortex, was related to the ability of individuals to infer the position of novel colors upon encountering them in a new rotation condition.

The OFC representation of mean spatial location on transfer trials specifically, was also correlated with behavioral evidence of successful transfer. We carried out an additional RSA analysis in this ROI only including transfer trials for the original colors in the first mini-block after a state change, with mean spatial position as the only regressor. While there was no significant effect for this regressor across participants (t(47) = 0.30, P = 0.7), the degree to which OFC distinguished between colors on original color transfer trials specifically, as measured by an RSA coefficient, was inversely correlated with participants’ mean normalized error on these trials (Supplementary Figure 11b; r(46) = -0.29, P = 0.04), further supporting a role for position coding in OFC during inference after rotations.

## Discussion

In this study, we developed a novel transfer learning task to examine whether grid codes rotate to facilitate the transfer of relational knowledge across different task states. We observed that individuals could leverage relational knowledge about the relative positions of target locations within one context to infer their locations in another – even in situations where the target locations had never been observed in the new context. In principle, such behavior could be supported by a cognitive grid-like code that undergoes rotations to remap associative knowledge into the relevant behavioral reference frame (Yu et al., 2021). Consistent with this notion, our fMRI analysis revealed hexagonally symmetric activity with respect to participants’ angle of movement in a set of brain regions that was very similar to those reported in past work (Bao et al., 2019; Constantinescu et al., 2016; Doeller et al., 2010), and that grid codes rotated into relevant behavioral reference frames in the rPPC. However, these parietal grid codes were not correlated with behavior and rotated too slowly to account for rapid transfer learning. Instead, we found that transfer behavior was more closely linked to representations of target identity and spatial location in the frontoparietal cortex and OFC, respectively.

### Neural grid representations that slowly rotate into cognitive reference frames

What could be the function of this slowly rotating grid-like code in the rPPC? The PPC has a crucial role in planning hand and eye movements in primates (Andersen et al., 1997). Human fMRI work has also frequently identified activity in the PPC during navigation tasks (Boccia et al., 2014), and other work has identified a hexagonally symmetrical signal for conceptual space in this region (Constantinescu et al., 2016; Doeller et al., 2010). This region receives weak input from the ERC in non-human primates (Munoz & Insausti, 2005), but could also receive signals from the ERC indirectly via the retrosplenial cortex (Kobayashi & Amaral, 2007) . Past work comparing the function of rodent PPC and ERC during navigation has argued that PPC activity is organized around egocentric actions rather than places, or context, suggesting a role in planning movement in a distinct and parallel frame of reference to that found in the ERC (Whitlock et al., 2012). The cognitive grid-like code observed here could be involved in facilitating some other behavior that we did not set out to characterize. For example, it could reflect the relative positions of the targets based on experience rather than their inferred positions after a state change. Another possibility is that the grid-like code observed here is not contributing anything meaningful to behavior but instead is simply an incidental by-product of how the brain parses two-dimensional task environments.

One surprising result from our study was that the bilateral ERC and several regions with significant hexagonal modulation did not maintain a consistent grid-like code that survived our cross-validation checks. Grid cells in the ERC are organized into modules that share the same phase (Stensola et al., 2012) , which is thought to allow for a grid-like code to be resolved by fMRI (Doeller et al., 2010). However, multiple such modules at different phases could co-exist at a sub-voxel level (Gu et al., 2018; Stensola et al., 2012), potentially interfering with our ability to resolve a coherent grid code. Electrophysiological studies in rodents have also revealed that the phase of grid cells is also not necessarily stable. While early work observed dramatic coherent rotations of the grid code occurring during changes of environment, more recent findings have shown that ERC neurons undergo remapping even in a stable environment (Low et al., 2021), and the phase of grid cells is less consistent over time than the relative phase of cells in different modules (Yoon et al., 2013). Notably, the coefficients measuring between-state consistency of the grid-like code were positively correlated in most of these ROIs across individuals. This result suggests that even though we could not observe a reliable grid-like code in most ROIs at the group level, the activity in these regions may reflect a common process related to spatial navigation. It is possible that different grid modules could be operating under a spatial or cognitive regime, and if these modules are intermixed at the voxel-level, we would not be able to resolve a coherent grid code. Future work recording directly from grid cells in a similar task could help resolve some of these ambiguities.

It is possible that the grid code may be operating differently in this predictive inference task than in typical spatial navigation settings. Other fMRI studies have been able to identify a grid-like code in the ERC and other regions during navigation of virtual physical and conceptual spaces (Bao et al., 2019; Doeller et al., 2010; Park et al., 2021). However, most of these tasks have involved a static environment with unambiguous, spatially fixed cues to indicate the current context. A notable exception is Julian and Doeller (2021), where participants carried out a spatial navigation task in two contexts with some trials where external cues rendered the context ambiguous. In that case, participants were trained on the two contexts separately, and during scanning could randomly enter either context in a given trial. However, unlike our task, the context in that experiment could be inferred from external cues, which may have facilitated the encoding of these contexts using grid-like codes with two distinct phases. Julian et al. (2018) also introduced a spatial rotation into their task and showed that the grid angle rotated in concert, but this rotation was visibly signaled by the layout of the display. It is possible that the grid code updates much more slowly for unsignaled changes in task state compared to those more dramatic, perceptual changes in state in other tasks. However, the slowness of the rotation of the grid-like code certainly raises questions about the mechanistic role these signals might serve in realistic rapidly changing real-world scenarios where the grid code has been suggested to play a role in making these kinds of fast inferences (Bellmund et al., 2018; Whittington et al., 2020; Yu et al., 2021).

### Neural identity and spatial codes predicted transfer across individuals

While we did not find evidence for the involvement of the grid code in transfer, we identified other neural representations that did correlate with this behavior across participants. We found that the OFC maintained a representation of the spatial layout of the target locations for shields in both states, and the strength of this representation was correlated with the degree to which individuals were able to infer novel shield color locations the first time observing them in a new rotation state. These findings are consistent with the notion that the OFC maintains a cognitive map of task space for inferring adaptive behaviors in novel situations (Wilson et al., 2014), and is consistent with the observation that OFC tracks shifting latent task states in a similar predictive inference task (Nassar et al., 2019), and maintains prospective goals during spatial navigation (Brown et al., 2016) . Interestingly, the OFC did not maintain a more abstract ‘invariant’ representation of the relative spatial positions of the shield colors that we anticipated would be involved in this transfer behavior, but instead encoded this information in concrete spatial terms. Representations of sensory specific information in the central OFC is thought to be involved in making inferences about concrete outcomes during decision-making about foods and odors (Howard et al., 2015; Howard & Kahnt, 2021). Similarly, the OFC may maintain concrete spatial information about target locations during navigation for inferring routes for goal-directed navigation behavior.

We also found that an indicator RDM for shield identify also correlated with transfer behavior. This RDM is relatively non-specific and may capture several different factors including shield color and learned stimulus-response rules for each color. As this regressor does not vary by state, it also captures between-state similarity in these representations. Given this non-specificity, it is not surprising that pattern activity in a large swath of frontal, parietal and visual cortex was correlated with this regressor. However, the strength of this representation in frontoparietal network was specifically correlated with transfer performance for the original shield colors. The network identified here is similar to that identified in many fMRI cognitive control tasks (Niendam et al., 2012) , thought to be involved in the flexible, context-dependent control of behavior (Badre & Nee, 2018; Miller & Cohen, 2001; Uddin, 2021) . This activity in the frontoparietal network could reflect a state-invariant rule-like representation of each target that facilitates transfer for original colors after a state change. These findings are consistent with the notion that different networks may maintain task information in distinct formats for the purpose of inference or efficient task execution (Vaidya & Badre, 2022). Namely, the spatial map in the OFC may facilitate inference about where to navigate for new shield colors that have not been tested before, while the frontoparietal network could facilitate rapid adjustments in behavior after state changes using an abstract representation of response rules for trained shield colors.

### Limitations

Some features of the task design place may limit interpretation of our results. This experiment involved navigating around an arena from a top-down perspective rather than using a first-person virtual reality display as is common in many other navigation experiments testing for a grid-like code using fMRI (Doeller et al., 2010; Julian & Doeller, 2021; Kunz et al., 2015; Raithel et al., 2023) . It is possible that the weakness of evidence for a grid-like code outside the rPPC could be the result of this design choice. However, other studies have found evidence for a grid code in visual spaces, using both fMRI in humans (Julian et al., 2018; Nau et al., 2018), and direct electrophysiological recordings in non-human primates (Killian et al., 2012). The layout of the task should also not be a factor of contention for theories of grid cell function that argue that these neurons are generally encoding information about the relational structure of the environment, and thus predict that this code is involved in navigation through other non-physical spaces (Behrens et al., 2018; Whittington et al., 2020) , as has been observed in several experiments (Bao et al., 2019; Park et al., 2021; Vigano et al., 2021). Those past experiments that tested for a grid-like code in visual space also used eye-tracking to measure movement in this space while we used the trajectory of participants’ shield movement. While it is possible that eye-movements might better measure participants’ plans during the simulation period, participants could also be planning future movements without changing fixation or through covert attention to target locations, in which case participants’ actual trajectory may better characterize the planned path in this task.

Our study was specifically designed to test a hypothesis about the function of the grid code in transfer behavior. For this reason, we only included two different rotation conditions in the fMRI session that would allow us to test predictions about a cognitive grid-like code and its relationship to transfer behavior. We also observed strong evidence of transfer behavior in the random rotation session, where participants demonstrated an ability to flexibly apply novel rotations to the shield positions. However, this condition was run as a behavioral follow-up study, limiting our ability to link neural representations to behavior within it. One promising avenue for future work would involve neuroimaging of this random rotation condition – allowing for direct and repeated measurement of the neural dynamics accompanying zero-shot inference.

### Conclusions

Here we showed that participants adaptively use relational knowledge to solve a spatial predictive inference task and transfer this knowledge to new scenarios. We observed that the rPPC maintains a cognitive grid-like code that rotates with behavioral reference frames in a manner that could enable it to transfer relational knowledge from one context to another. However, we found no evidence that this signal relates to transfer behavior either within or across individuals and demonstrated that it rotated too slowly to facilitate rapid relational inferences. Other representations of task relevant information about target identity and spatial location do appear to be involved in transfer behavior, arguing for a different set of neural and cognitive mechanisms. Our work sheds light on the representations facilitating rapid transfer learning in humans and provides important constraints shaping our understanding of the role that grid representations play in cognition.

## Methods

### Participants

Sixty-five participants were recruited for this three-session behavior/MRI study. Four participants were excluded due to coding errors in the behavioral task. Seven participants dropped out after the first session either voluntarily or due to MRI exclusion criteria, and six participants were excluded due to either excessive motion during scanning (translational movement in any direction exceeding 1 voxel during one or more runs) or falling asleep during the scanning session. Forty-eight participants (25 female, mean age = 23.9 years, SD = 4.9 years) were included in the study. Two participants did not have scanner data for the last run of the fMRI session due to time constraints or experimenter error but are included in the study. One participant did not complete the third session of the study, but their data for the first two sessions are included in all other analyses. All participants gave their written informed consent to participate in this study, as approved by the Human Research Protections Office at Brown University, and were compensated for their participation.

### Experimental design

#### Training session (Session 1)

Participants completed a predictive inference task that required using a joystick to navigate their “shield” a location on a circular arena (“planet”) where an “alien attack” was expected to occur. Predictions were possible because the shield color was related to the attack location allowing participants to learn color-location associations through experience.

Each trial was composed of three parts: simulation, response, and feedback. Each trial began with a simulation period, the duration of this period was randomly drawn from a uniform distribution to be 2-6 seconds in 1 second intervals. During this period, the shield was initialized at a random location on the screen. Participants were instructed to imagine moving the shield to the location they want to during this period. After the simulation period, participants were queried for a response. The start of the response period was signaled by the grey outline of the planet turning red, upon which participants had 3 seconds to move the shield (at a constant speed) to its predicted location using a joystick. Immediately following the response period, the outcome (the “bomb”) appeared as a dot for 1 second, which was colored red if the participant’s shield missed the location of an alien attack, and green if the shield overlapped with the bomb.

Participants were required to learn the most probable attack locations associated with each of five different shield colors. Each color was associated with a target location corresponding to the center of attacks for that color. Target locations were constrained such that their radial component was greater than 20%, but less than 80%, of the planet radius, ensuring that angular error measurements would be meaningful and minimizing edge effects. Trial outcomes were drawn from a two-dimensional Gaussian distribution centered on the target location for the current shield color and with a relatively small variance (Std = 4% of the planet radius). Participants were informed regarded the variance around the target location and were instructed to try to block as many attacks as possible by estimating the true underlying target location.

The task was composed of sets of interleaved trials interrupted at fixed points by occasional state changes. Shield colors were interleaved in mini-blocks that included one presentation of each of the five shield colors in pseudorandom order. A state change would occur after eight such mini-blocks, at which time target locations associated with each color were rotated by 90° with respect to the circular arena. Participants performed another state-block of the same length (8 mini-blocks) and were then allowed a break. Prior to the start of the subsequent block, another state transition would occur, this time rotating all color targets counter clockwise (−90° with respect to the circular arena) back into their original positions. Participants completed eight state-blocks of trials in total (320 trials) and were incentivized with a bonus payment that depended on how many bombs they “caught”.

#### Novel colors inference session (Session 2)

On session 2, participants completed a similar behavioral task while they underwent fMRI scanning. The shield color -attack location associations of the five shields they had learned about in Day 1 were carried over to this session. There were now six mini-blocks per state-block in this session, and participants experienced alternating rotations between 0° and 90° as in the first session.

In addition, the participants learnt about eight additional color-attack location associations (referred to as “novel color trials”). The novel color trials would only be presented after the sequence of the original five colors were presented in each stimulus repetition. There were two novel color trials per mini-block. Participants would be introduced to a novel color and learn about its location association during one state-block, and performance on the first instance of the same color (thus prior to any feedback) on the subsequent rotated state-block is taken as evidence for transfer. There were three types of novel color trials: for the first type, the novel color was presented for the first time in the original state-blocks, and the inferential test is in the subsequent rotated state-blocks. The second type presented the novel color for the first time in the rotated state-blocks, and the inferential test was in the subsequent original state-blocks. Finally, there was one trial type where the novel color was presented in the original state-block at the beginning of the session, and the inferential test in the rotated state-block was at the end of the experiment. This last trial type was not included in any analyses, as the long time between training and transfer made these trials very different from other novel color trials. Participants completed eight state-blocks in total (332 trials).

#### Novel rotations inference session (Session 3)

On the third session, participants again encountered the same color-attack location associations (at the 0° rotation positions) of the original five colors in the first block of the experiment. After the first state-block, the attack locations corresponding to the same colors were rotated by one of 23 randomly selected potential configurations between 15-345° along 15° intervals, rather than the +90 and -90° rotations that they had experienced previously. In each subsequent state change, the target locations rotated again to another of these configurations, randomly selected without replacement. Participants experienced eight mini-blocks per state-block, and 8 state-blocks in total (with 7 blocks being novel rotation state-blocks, 320 trials total).

#### Behavioral analysis

To analyzed participants’ responses, we calculated a measure of normalized angular error

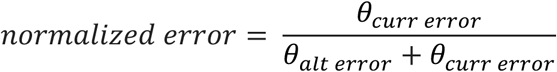

Where *θ_curr error_* is the angular distance between the terminal shield position for the participant’s response and the mean target position for the shield in the current state, and *θ_alt error_* is the angular distance between the terminal shield position and the mean target position for the shield in the alternate state. All angles were referenced to the center of the arena. This measure normalizes participants angular error to a scale from 0 to 1, where a number closer to 0 signifies a response more appropriate for the current state, and a number closer to 1 signifies a response more appropriate for the alternative state, and a number closer to 0.5 signifies a random response equidistant between the target locations for the two states.

In each session, we analyzed how this normalized error changed over multiple time scales with multiple linear regression analyses carried out on individual participants and group-level tests carried out through two-tailed one-sample t-tests on regression coefficients across participants. We conducted two tests, one focused on the first mini-block after a state change with regressors for trial number, block number and their interaction, and a second regression model at the larger scale of mini-blocks, with regressors for mini-block number, block number and their interaction.

To compare participants’ task performance with different neural measures, we calculated a measure of error on transfer trials for the original colors (i.e. mean error on trials 2-5 in the first mini-block after a state change), and for the novel colors specifically in session 2. As these values were on a restricted 0-1 scale, the measures were logit transformed before being submitted to Pearson correlations with neural measures. However, in all accompanying scatterplots we show the original untransformed data for better interpretability.

### Transfer error analysis

We sought to test whether individual participants’ absolute angular error on transfer trials was less than what would be expected based on a random guessing strategy. To this end, we simulated 1000 means for each participant with a random response angle on each trial. We then sorted these responses and took the 97.5 and 2.5 percentiles of these errors across simulated responses in each simulated participant. This provided us with the 95% confidence interval for the angular error for a random response strategy which we could compare to our participants’ angular error on transfer trials to determine whether their responses were outside this range.

### fMRI acquisition

Whole-brain imaging was collected using a Siemens 3T Magnetom Prisma system with a 64-channel head coil. In each fMRI session, a high resolution T1 weighted MPRAGE image was acquired for visualization (repetition time (TR), 1900 ms; echo time (TE), 3.02 ms; flip angle, 9°; 160 sagittal slices; 1 × 1 × 1 mm voxels). Functional volumes were acquired using a gradient-echo echo planar sequence (TR, 2000 ms; TE, 25 ms; flip angle 90°; 40 interleaved axial slices tilted approximately 30° from the AC-PC plane; 3 × 3 × 3 mm voxels). Functional data were acquired over four runs. Each run lasted 11.53 min on average (346 acquisitions). Soft padding was used to restrict head motion throughout the experiment. Stimuli were presented on a 32-inch monitor at the back of the bore of the magnet, and participants viewed the screen through a mirror attached to the head coil. Participants used a joystick pad to interact with the experiment.

### fMRI preprocessing

The fMRI data in this manuscript is preprocessed using *fMRIPrep,* a Nipype based pipeline (Estaban et al. 2019). The description below is copied from the *fMRIPrep* boilerplate text, with parts not used for analyses removed.

### Anatomical data preprocessing

The T1-weighted (T1w) image was corrected for intensity non-uniformity (INU) with N4BiasFieldCorrection (Tustison et al., 2010), distributed with ANTs 2.2.0 (Avants et al., 2008), and used as T1w-reference throughout the workflow. The T1w-reference was then skull-stripped with a *Nipype* implementation of the antsBrainExtraction.sh workflow (from ANTs), using OASIS30ANTs as target template. Brain tissue segmentation of cerebrospinal fluid (CSF), white-matter (WM) and gray-matter (GM) was performed on the brain-extracted T1w using fast (FSL 5.0.9, Zhang et al. (2001)). Volume-based spatial normalization to one standard space (MNI152NLin2009cAsym) was performed through nonlinear registration with antsRegistration (ANTs 2.2.0), using brain-extracted versions of both T1w reference and the T1w template. The following template was selected for spatial normalization: *ICBM 152 Nonlinear Asymmetrical template version 2009c* (Fonov et al., 2009); TemplateFlow ID: MNI152NLin2009cAsym].

### Functional data preprocessing

For each of the 4 BOLD runs found per subject (across all tasks and sessions), the following preprocessing was performed. First, a reference volume and its skull-stripped version were generated using a custom methodology of *fMRIPrep*. Susceptibility distortion correction (SDC) was omitted. The BOLD reference was then co-registered to the T1w reference using flirt (FSL 5.0.9,Jenkinson and Smith (2001)) with the boundary-based registration (Greve and Fischl 2009) cost-function. Co-registration was configured with nine degrees of freedom to account for distortions remaining in the BOLD reference. Head-motion parameters with respect to the BOLD reference (transformation matrices, and six corresponding rotation and translation parameters) are estimated before any spatiotemporal filtering using mcflirt (FSL 5.0.9, Jenkinson et al. (2002)). BOLD runs were slice-time corrected using 3dTshift from AFNI 20160207 (Cox & Hyde, 1997). The BOLD time-series (including slice-timing correction when applied) were resampled onto their original, native space by applying the transforms to correct for head-motion. These resampled BOLD time-series will be referred to as *preprocessed BOLD in original space*, or just *preprocessed BOLD*. The BOLD time-series were resampled into standard space, generating a *preprocessed BOLD run in [‘MNI152NLin2009cAsym’] space*. First, a reference volume and its skull-stripped version were generated using a custom methodology of *fMRIPrep*. Several confounding time-series were calculated based on the *preprocessed BOLD*: framewise displacement (FD), DVARS and three region-wise global signals. FD and DVARS are calculated for each functional run, both using their implementations in *Nipype* (following the definitions by Power et al. (2014). The three global signals are extracted within the CSF, the WM, and the whole-brain masks. All resamplings can be performed with *a single interpolation step* by composing all the pertinent transformations (i.e. head-motion transform matrices, susceptibility distortion correction when available, and co-registrations to anatomical and output spaces). Gridded (volumetric) resamplings were performed using antsApplyTransforms (ANTs), configured with Lanczos interpolation to minimize the smoothing effects of other kernels (Lanczos, 1964).

### General linear models

fMRI data were analyzed with general linear models (GLMs) using SPM12. Each of these models included three cosine components capturing low-frequency slow fluctuations in the signal (from 0.5-2 periods of each run), six regressors for translational and rotational head motion and their temporal derivatives, mean signal from the CSF and WM and individual regressors for frames labeled as motion outliers. The threshold for motion outliers was changed to 2.5 standard deviations of the standardized DVARS above the mean standardized DVARS or 0.5 mm framewise displacement, except where this threshold was below 1.5 standardized DVARS, where we reverted to the fMRIprep defaults (1.5 standardized DVARS or 0.5 mm framewise displacement). In all GLM models, separate regressors were included for correct, incorrect and missed responses. These regressors were modeled boxcars of the same duration as the trial simulation period. Trials were categorized as ‘correct’ when the angular error was under 45 degrees (i.e. less than half of the angular distance between the two states in the task on Day 1 and 2), categorized as incorrect if the angular error was above this threshold, or categorized as a missed responses when the shield was moved by less than 10 pixels independent of the angular error. Additional boxcar regressors were also included for outcomes and responses, with the durations matching the length of these periods (1 s and 3 s, respectively).

### Hexagonal modulation analysis

A whole brain analysis was conducted to test where activity was consistent with a hexagonally symmetric signal. We adopted the approach taken by (Constantinescu et al., 2016), including two parametric modulators for each simulation period regressor corresponding to the sine and cosine of the angle of travel in each trial (*θ*), with a periodicity of 60 degrees, that is sin(6*θ*) and cos(6*θ*). Separate regressors and parametric modulators were included for correct and erroneous responses in both the original and rotated states. All runs were concatenated together to provide the maximum number of trials in which to estimate beta coefficients for each of these modulators. We then tested where brain activity showed strong hexagonal modulation independent of state using an F-test on the linear combination of the beta coefficients for the sin(6*θ*) and cos(6*θ*) regressors for correct responses in both states. These F-statistics were then transformed to Z-statistics using an asymptotic transformation (http://www.fmrib.ox.ac.uk/analysis/techrep/tr00mj1/tr00mj1/), and the distribution of Z-statistics was tested against zero using a one-sample t-test at the whole-brain level, corrected for multiple comparisons (see below).

### Between-state grid-like code tests

To test if regions that showed evidence of hexagonal modulation of the BOLD signal at the whole-brain level demonstrated a reliable cognitive or spatial grid-like code, we carried out between-state cross-validation tests in ROIs centered on effects from this whole-brain analysis (see below). In each ROI, we used the same GLM for the hexagonal modulation analysis to estimate beta coefficients for the sine and cosine parametric modulators in each state. We then calculated a grid angle *φ* for each ROI in a given state X using the method developed by Doeller et al. (2010), where *φ*_state X_ = [*arctan(β_sin-state X_/β_cos-state X_*)]/6. We then used this grid angle from state X to create sixfold parametric modulators for the simulation period regressor reflecting the expected BOLD response in the other state Y with *cos(6[θ_t_-φ*_state X_]), where *θ_t_* is the angle of travel in each trial. We replaced the sine and cosine parametric modulators in the GLM used for the hexagonal modulation analysis with these parametric modulators such that the modulator for trials from state Y reflected the predictions based on the grid angle for state Xand vice-versa.

To examine if these ROIs demonstrated either a reliable cognitive or spatial grid-like code between states, we tested if the mean coefficients for the sixfold cosine parametric modulators for the original and rotated states were significantly above or below zero using a two-tailed, one-sample t-test across participants. For a cognitive grid-like code where the angle is fixed to the relative position of the target locations for the shield colors, we expected that the BOLD response in state Y would be anti-phase to the activity predicted by a grid angle fit from state X and vice-versa and therefore these coefficients should be more negative. However, for a spatial grid-like code fixed to the real physical layout of the display, we would expect the predicted responses in states X and Y to be in-phase, and thus expected these coefficients to be more positive.

### Regions-of-interest

For all analyses on ROIs, a spherical mask was created centered on the peak MNI coordinates of each effect of interest with a 10 mm radius using the MarsBar toolbox (Brett et al., 2002). This mask was isolated to gray matter voxels by removing voxels where the SPM12 gray matter tissue probability map was below 0.5.

### Grid-like code reliability and specificity tests

To test the within-state reliability of the grid-like code observed in the rPPC, we created a new GLM where runs were not concatenated. This allowed us to estimate separate parametric modulators for the sine and cosine of the angle of travel in each state and each run. Using the same methods described above, we calculated a grid angle for each state and run (*φ*_stateX,run1_, *φ*_stateX,run2_, etc.) and then carried out a leave-one-out cross-validation procedure where we calculated the mean grid angle for each state X for all but one run (e.g. *φ*_state X, runs 1..n-1_), and used this to grid angle to generate a parametric modulator for the same state X in the held out run (e.g. run *n*). We repeated this process for all runs in both states, and took the average of the coefficients for the modulators across runs and states in each participant. We then compared these coefficients against zero with a one-sample, right-tailed t-test.

To test the specificity of this grid-like code to a sixfold modulator, we also carried out the same between-and within-state tests described above, only changing the number of folds for the rotational symmetry to 4, 5, 7 or 8 (90°, 72°, 51.4° and 45° periodicities).

### Trial-wise analysis

Trial-wise measures of the grid-like code were calculated using a method adapted from Julian and Doeller (2021) to derive estimates of which state (i.e. the rotated or original state) the signal in this region was consistent with at a trial-wise level. Using the grid angles for each state from the between-state cross-validation analysis, we generated single-trial regressors for each trial using the same sixfold cosine function used to generate parametric modulators for the between-and within-state cross-validation analyses above, such that the sign and amplitude of the regressor for a single trial depended on the trial angle of shield movement and the grid angle from the alternate state. We then estimated Pearson correlation coefficients between these regressors and the residualized BOLD signal in this ROI after removing linear effects for trial and movement regressors. As with the between-state cross-validation analysis, because the grid angle used to generate these regressors was from the alternate state, the sign of these coefficients could be interpreted as evidence of the persistence of the grid code from the alternate state (positive), or a rotation of 90° consistent with a cognitive grid-like code (negative). These correlation coefficients were then Fisher transformed before being submitted to statistical tests.

We tested the dynamics of this trial-wise measure and its relationship with behavior using multiple linear regression analyses in individual participants and one-sample t-tests on participant-level coefficients. We first carried out a regression model with trial-wise coefficients as the dependent variable and the logit transformed normalized error as the dependent variable. We also conducted a second test including the number of mini-blocks after state change as a covariate (as this was strongly predictive of changes in participants’ error). To test how these trial-wise coefficients changed over multiple time courses, we carried out two linear regression analyses testing the relationship between these coefficients and trials in the first mini-block immediately after a state-change, and with mini-blocks after a state change.

### Representational similarity analysis

To identify correlates of task variables that could putatively support transfer behavior, we carried out an RSA analysis on a trial-by-trial RDM to examine. We used a single-trial GLM approach to estimate a t-statistic map for the simulation phase in each trial (Mumford et al., 2012). For each trial, this GLM had a single regressor for the simulation period of the trial of interest and regressors for all other simulation periods, all outcomes, all responses, and the same nuisance regressors as were used for the GLMs used in the grid code analyses above. We conducted a whole-brain searchlight analysis with a circular searchlight that had a 9 mm radius. In each pass of this searchlight, we calculated trial-by-trial neural RDMs using the Pearson correlation distance for all voxels in this searchlight.

The lower triangle of this neural RDM was vectorized and compared to vectorized lower triangle of multiple hypothesis RDMs and nuisance covariates using a multiple linear regression analysis. Among the regressors of interest were RDMs that captured trial-wise measures of the following:

1. the Euclidean distances of the mean spatial position of the target location for each shield color
2. the Euclidean distances between the mean spatial position of the target location for each shield color that was invariant across both states
3. an indicator for shield identity where trials with the same shield color were more similar to each other and equally dissimilar from all other trials
4. an indicator for transfer trials (i.e. trials from the first mini-block after a state change), where these trials were similar to each other and equally distant from all other trials

Covariate regressors included RDMs that captured the following:

1. An indicator for the scanner run
2. An indicator for the state-block in each run
3. An indicator for the current task state (i.e. whether trials were in the original or rotated states)
4. An indicator for shield identity, constrained to within-state comparisons
5. Trial-wise absolute differences in the angular change of the participant’s response from the last presentation of the color
6. Perceptual distances between shield colors in each trial based on the CIELAB color space
7. Shield movement, measured as the Euclidean distance between the start and end shield positions on each trial
8. Euclidean distance for the shield starting positions in each trial
9. Euclidean distances for the shield end position in each trial
10. Lag regressors capturing temporal autocorrelation between off-diagonal pairs of trials from trial t+1 to t+16

In each pass of this searchlight, beta coefficients for all regressors were calculated at each step and averaged over searchlight passes for all voxels included in the searchlight to compute fixed-effects in a first-level analysis. For second-level tests, coefficients from each of these regressors at each voxel were compared against zero using a t-test. Statistical significance for the resulting t-statistic maps was then assessed using non-parametric permutation tests to correct for multiple comparisons (see below).

To test where correlates of these RDMs were related to behavioral performance, we carried out additional second-level tests where behavioral measures were included as a covariate of interest in the second-level model. In these cases, we only applied a mask so that this test was only conducted in voxels where there were significant effects for the RDM at a whole-brain, cluster corrected level.

### Second-level whole brain tests

For all whole brain statistical tests, t-statistic maps were thresholded at a cluster defining threshold of p<0.001 uncorrected, and a cluster extent threshold of P < 0.05, corrected for multiple comparisons using non-parametric statistics. The cluster extent threshold (k) was defined using non-parametric permutation tests (10,000 permutations) where the sign for half of participants’ contrasts was randomly flipped in each permutation to generate a null distribution based on the suprathreshold maximum cluster statistics. The cluster extent was defined as the 95^th^ percentile of this null distribution.

## Acknowledgements

We would like to thank members of the Learning, Memory and Decision Lab for their constructive feedback on this work and to Haoxue Fan for providing comments on the manuscript. Additional thanks to Jeroen van Baar for assistance implementing the preprocessing pipeline, and Nir Moneta for helpful analysis suggestions. This work was supported by a COBRE Center for Central Nervous System Function Pilot Award (1P20 GM103645), and an award from the National Institute on Mental Health (R01MH126971). This research was supported in part by the Intramural Research Program of the National Institutes of Health (NIH) (ZIA DA000642). The contributions of the NIH author are considered Works of the United States Government. The findings and conclusions presented in this paper are those of the author and do not necessarily reflect the views of the NIH or the U.S. Department of Health and Human Services.

---

**Supplementary Figure 1.**
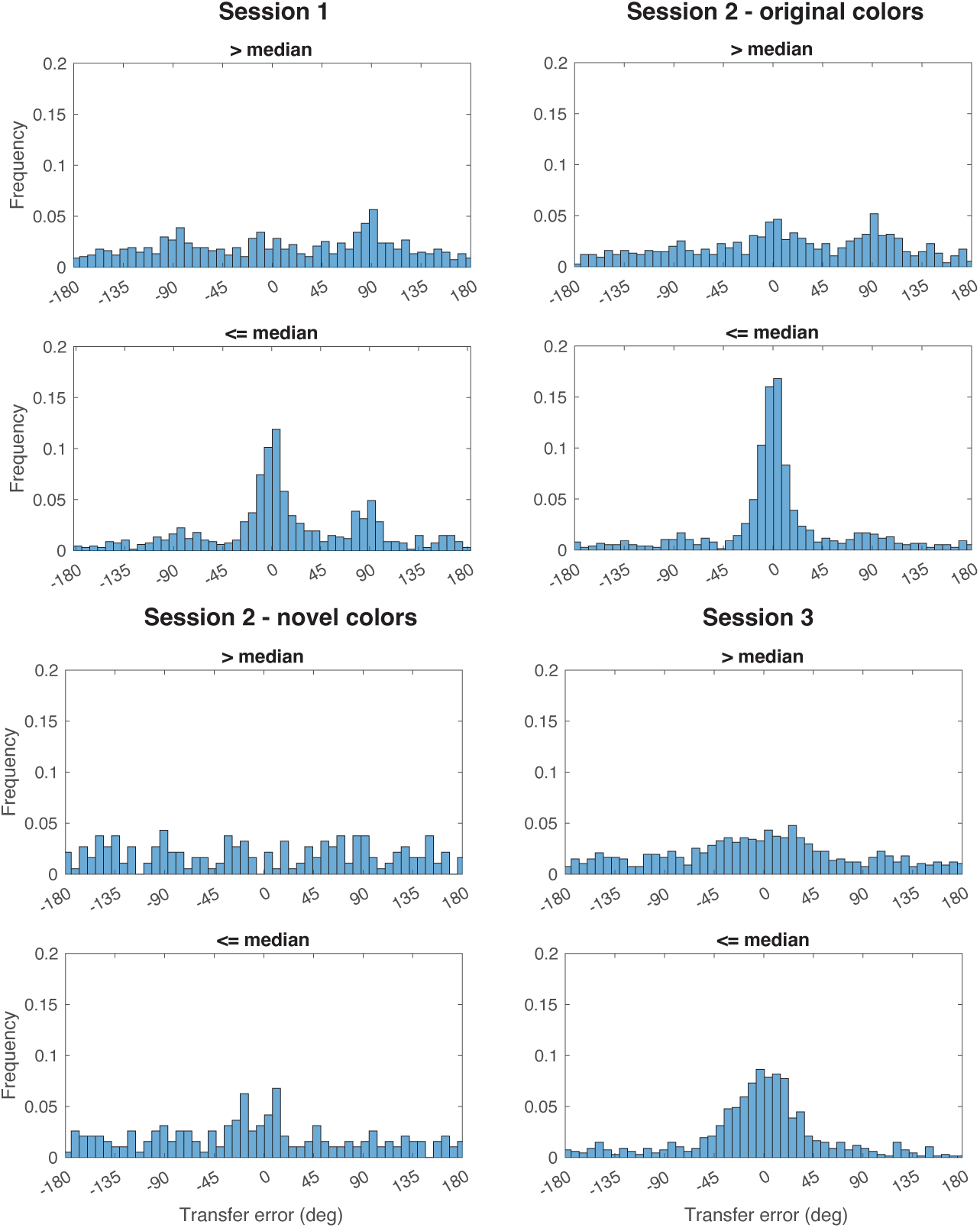
Histograms showing angular error on transfer trials measured in degrees across sessions and conditions. Participants have been separated based on a median split of the average transfer error in each condition. Note the peak centered on zero degrees for the top half of participants (i.e. <= median transfer error) in the novel color condition in session 2.

**Supplementary Figure 2.**
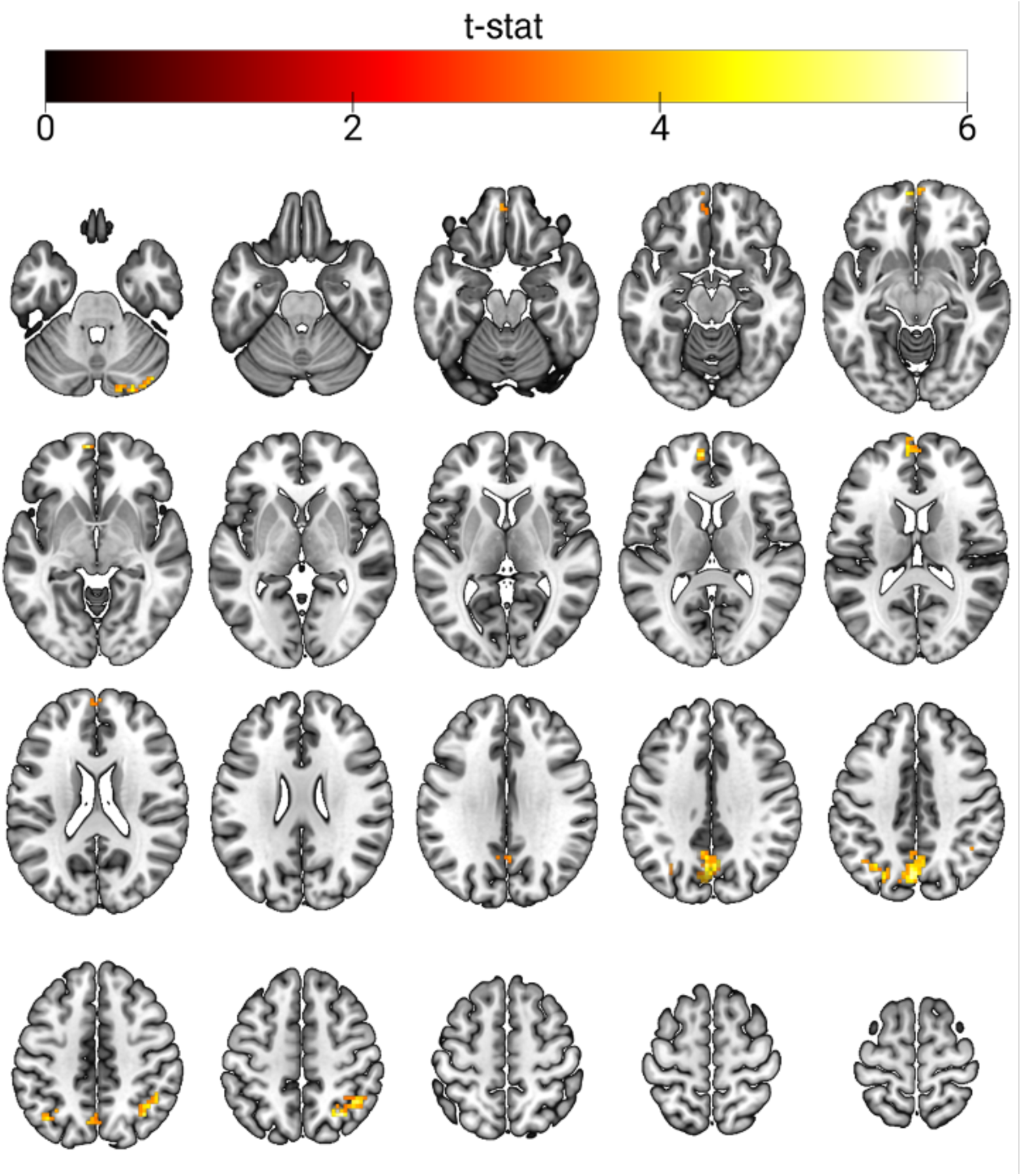
Statistical map from whole brain analysis identifying regions responding to a hexagonally symmetrical signal during the response period. Results are displayed at a cluster forming threshold of P<0.001 and corrected for multiple comparisons with permutation tests for defining a cluster extent threshold at P<0.05 (K = 52).

**Supplementary Figure 3.**
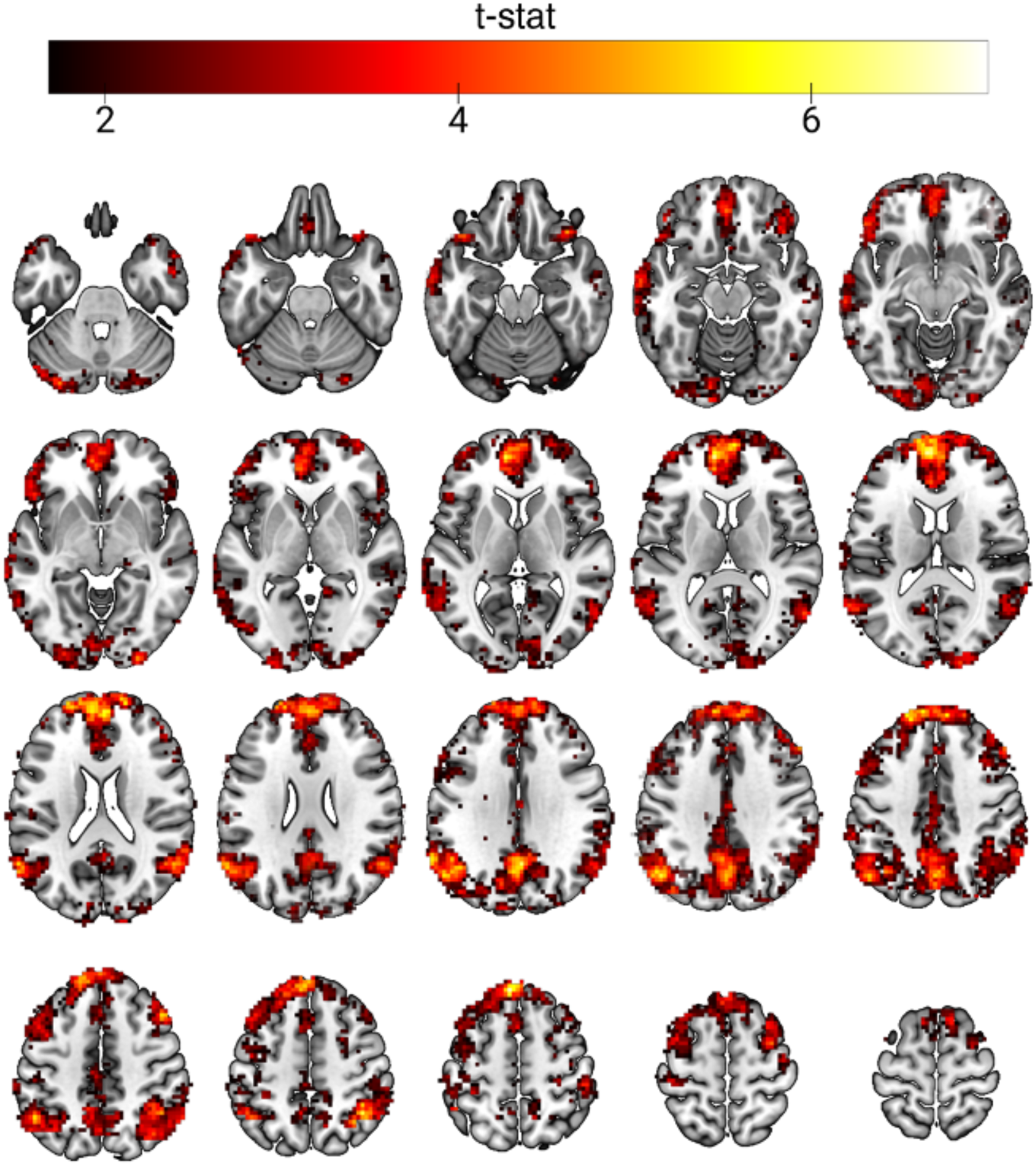
Statistical map from whole brain analysis identifying regions responding to a hexagonally symmetrical signal at an uncorrected threshold of P < 0.05.

**Supplementary Figure 4.**
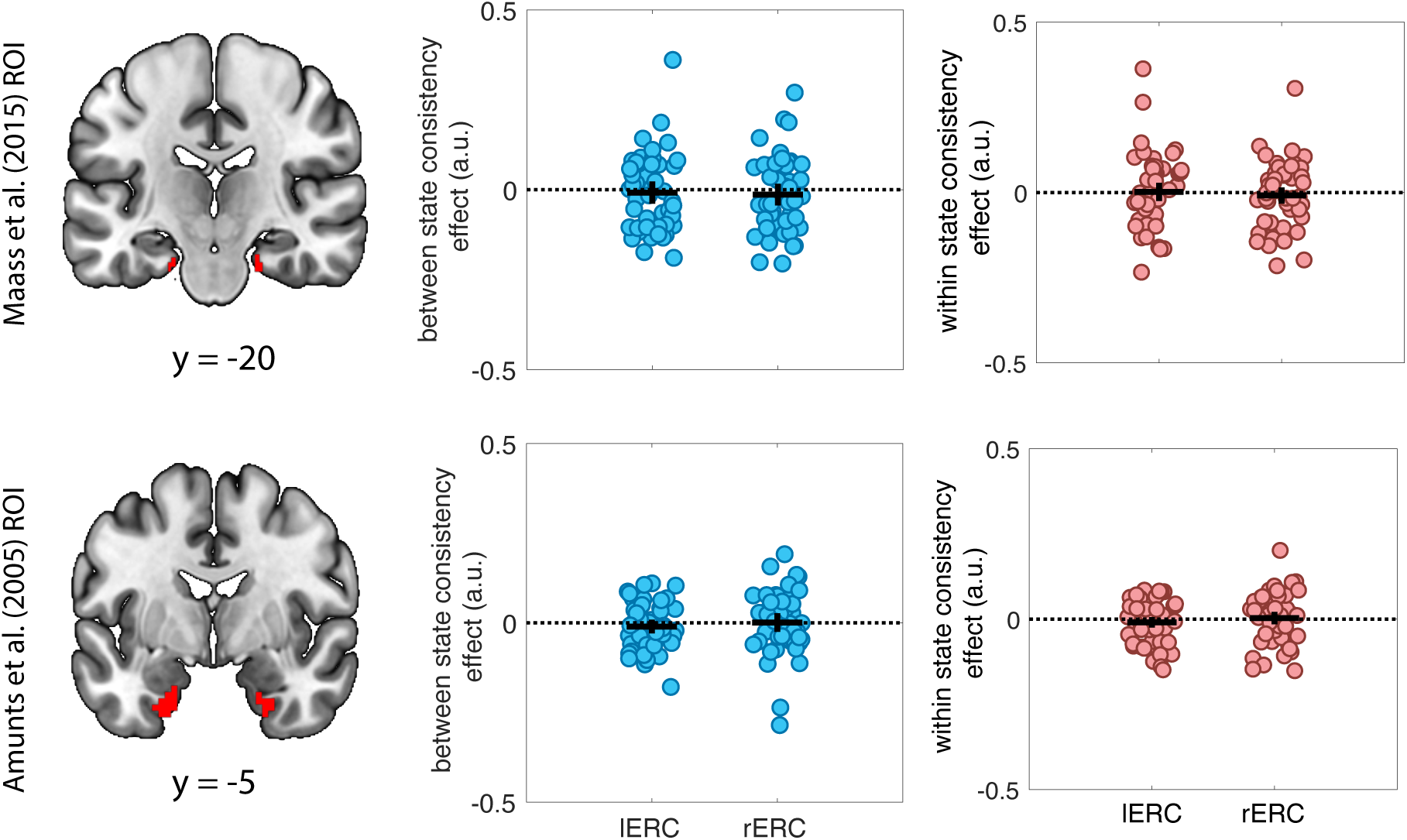
Tests for between- and within-state grid angle consistency in two pairs of *a priori* entorhinal cortex ROIs in the left and right hemisphere (lERC, rERC) in which other fMRI studies that have identified grid-like codes (Bao et al., 2019; Park et al., 2021; Raithel et al., 2023). These ROIs are shown on the left-hand side. Each circle represents a participant. Horizontal bars represent the mean, vertical bars represent the 95% CIi. There were no significant results in either set of ROIs.

**Supplementary Figure 5.**
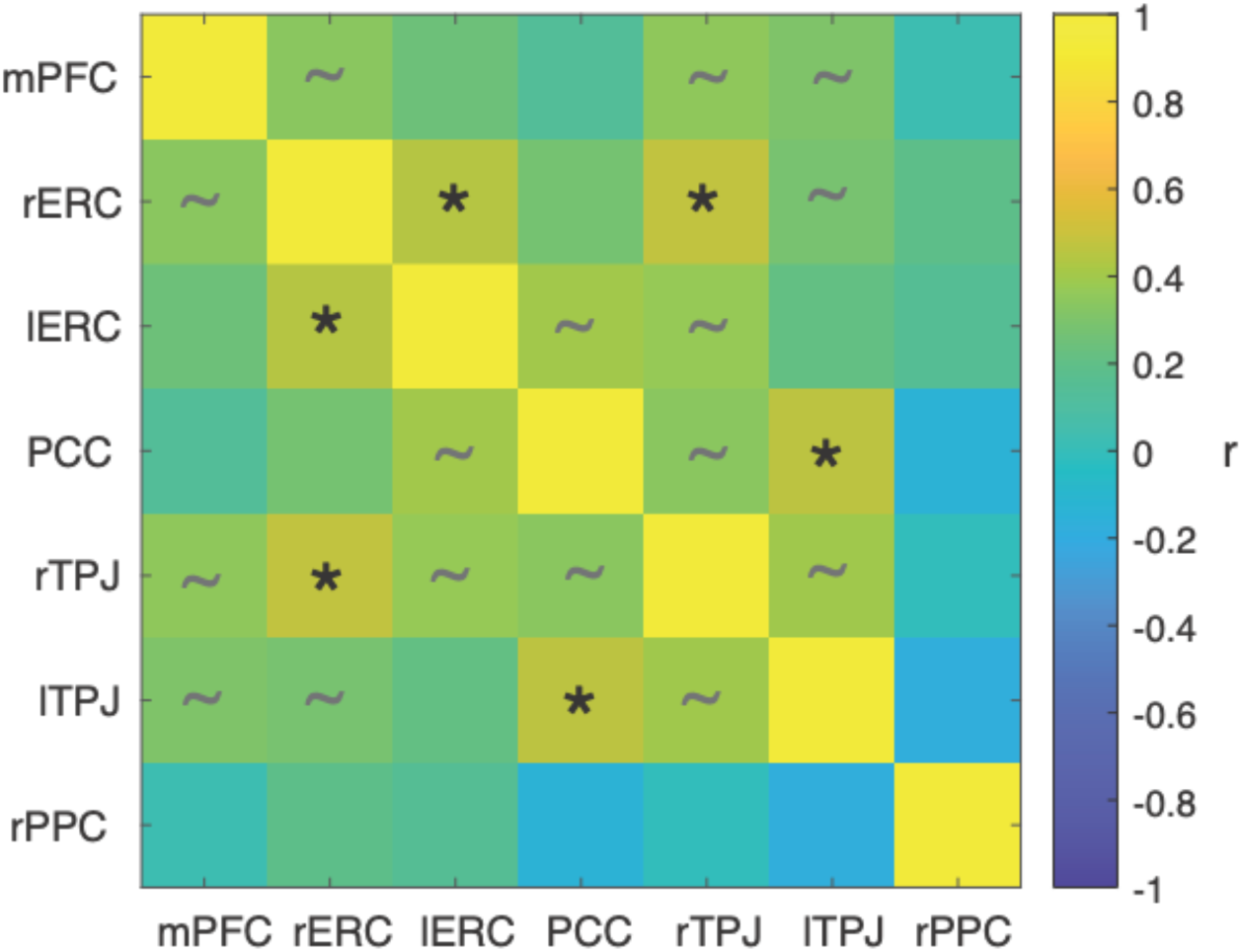
Pearson correlation matrix for between-state grid coefficients for different regions of interest. We identified significant positive correlations between these coefficients in the right and left ERC, and between the right temporoparietal junction (TPJ) and right ERC, and between the posterior cingulate cortex and left TPJ (r’s(46) ≥ 0.44, P < 0.05, corrected for multiple comparisons). Coefficients in other regions identified in the hexagonal symmetry analysis were also weakly positively correlated, though these correlations did not survive multiple comparisons correction. Notably, these correlations were weakest or trended in a negative direction between these other ROIs and the rPPC, which was the only brain region with grid representations that had a consistent phase relationship across rotation states. Medial prefrontal cortex (mPFC), entorhinal cortex (ERC), posterior cingulate cortex (PCC), tempro-parietal junction (TPJ), posterior parietal cortex (PPC) r = right, l = left. * P < 0.05, corrected for multiple comparisons, ∼ P < 0.05 uncorrected.

**Supplementary Figure 6.**
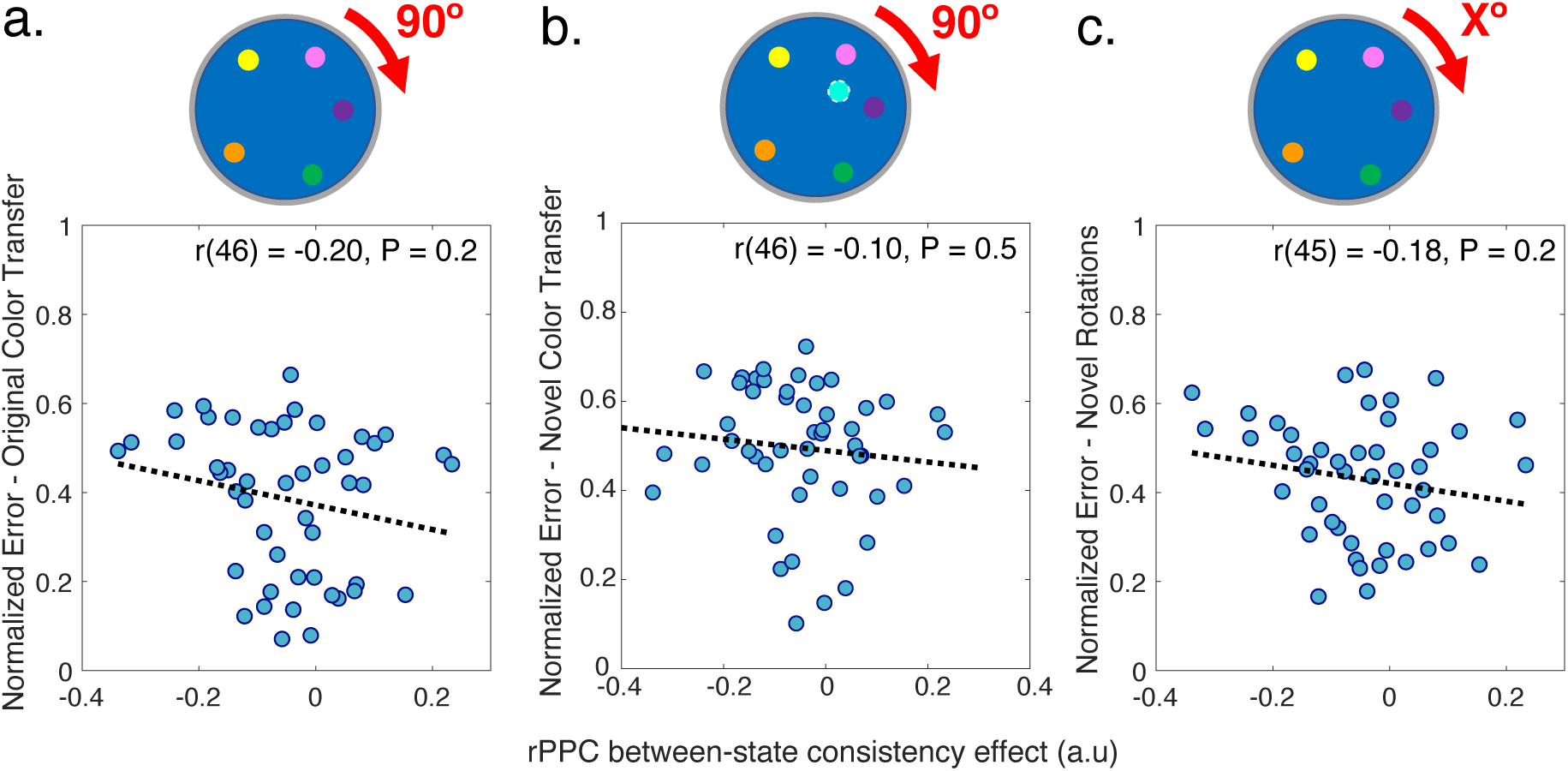
Scatterplots between right posterior parietal cortex (rPPC) between-state grid-like code consistency and normalized error on transfer trials for original colors (**a**) and novel colors (**b**) in session 2, and novel rotations in session 3 (**c**).

**Supplementary Figure 7.**
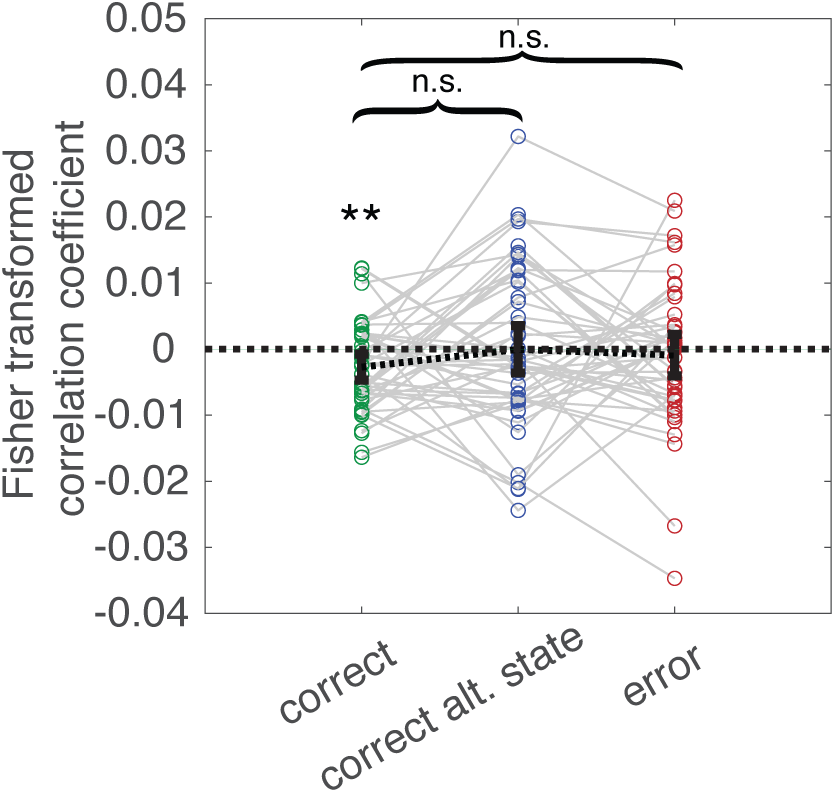
Trial-wise coefficients for between state grid-like code phase consistency organized by trial type. We expected that if this cognitive grid-like code in rPPC reflects participants belief about the task state, then the grid angles should be anti-phase between states on trials where they respond correctly given the current state (i.e. more negative coefficients), and would be in-phase between states in trials where participants responded as though they were in the alternate state. On correct trials, where participants were within 45 degrees of the target, there was a significant negative tendency for these coefficients, two-tailed one-sample t-test: t(47) = 2.76, P = 0.008, d = 0.40), as expected. However, coefficients on trials that were correct for the alternate state (i.e. within 45 degrees of the target location in the alternate state) were not significantly different from zero (Figure 5a; t(47) = -0.02, P = 0.9, d = 0.003). There were also no significant differences between coefficients on correct trials and trials correct for the alternate state (t(47) = 1.36, P = 0.18, d = 0.20), or between correct trials and other kinds of error trials (t(47) = 0.96, P = 0.34, d = 0.14).

**Supplementary Figure 8.**
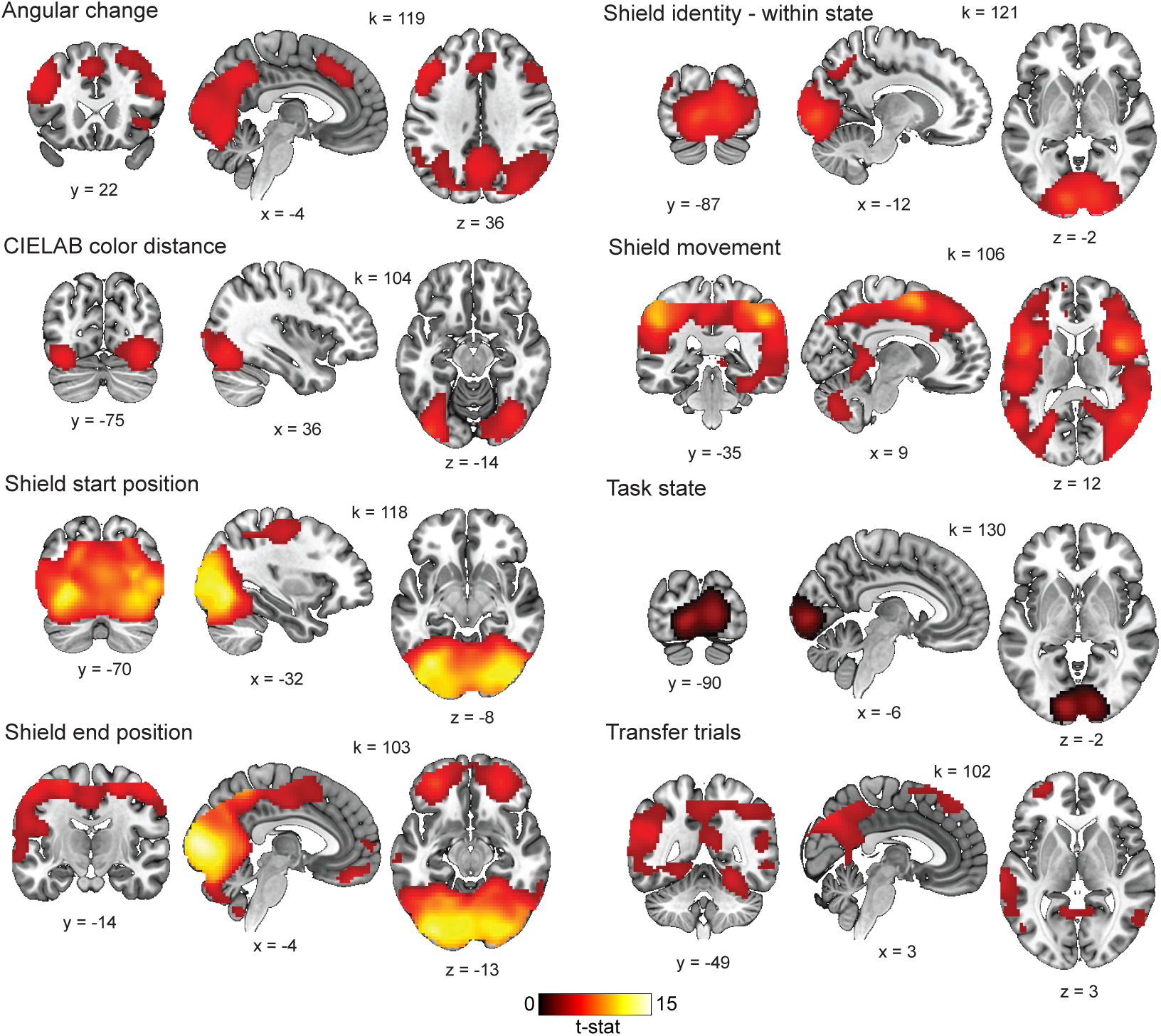
Results of whole-brain representational similarity searchlight analysis for all task and behavior related regressors with statistical effects that survived multiple comparisons correction. All statistical maps were defined with a cluster forming threshold of p<0.001 and corrected for multiple comparisons with permutation tests for defining a cluster extent threshold at p<0.05. Cluster extent threshold for each contrast is given by the value of k. Covariate analyses restricted to voxels that showed a significant effect for the transfer RDM at a whole-brain cluster corrected threshold did not reveal any significant correlations with normalized error on transfer trials for original or novel colors. We also did not find any regions with a significant representation of the invariant positions of each shield color, even at a more liberal threshold of P < 0.01.

**Supplementary Figure 9.**
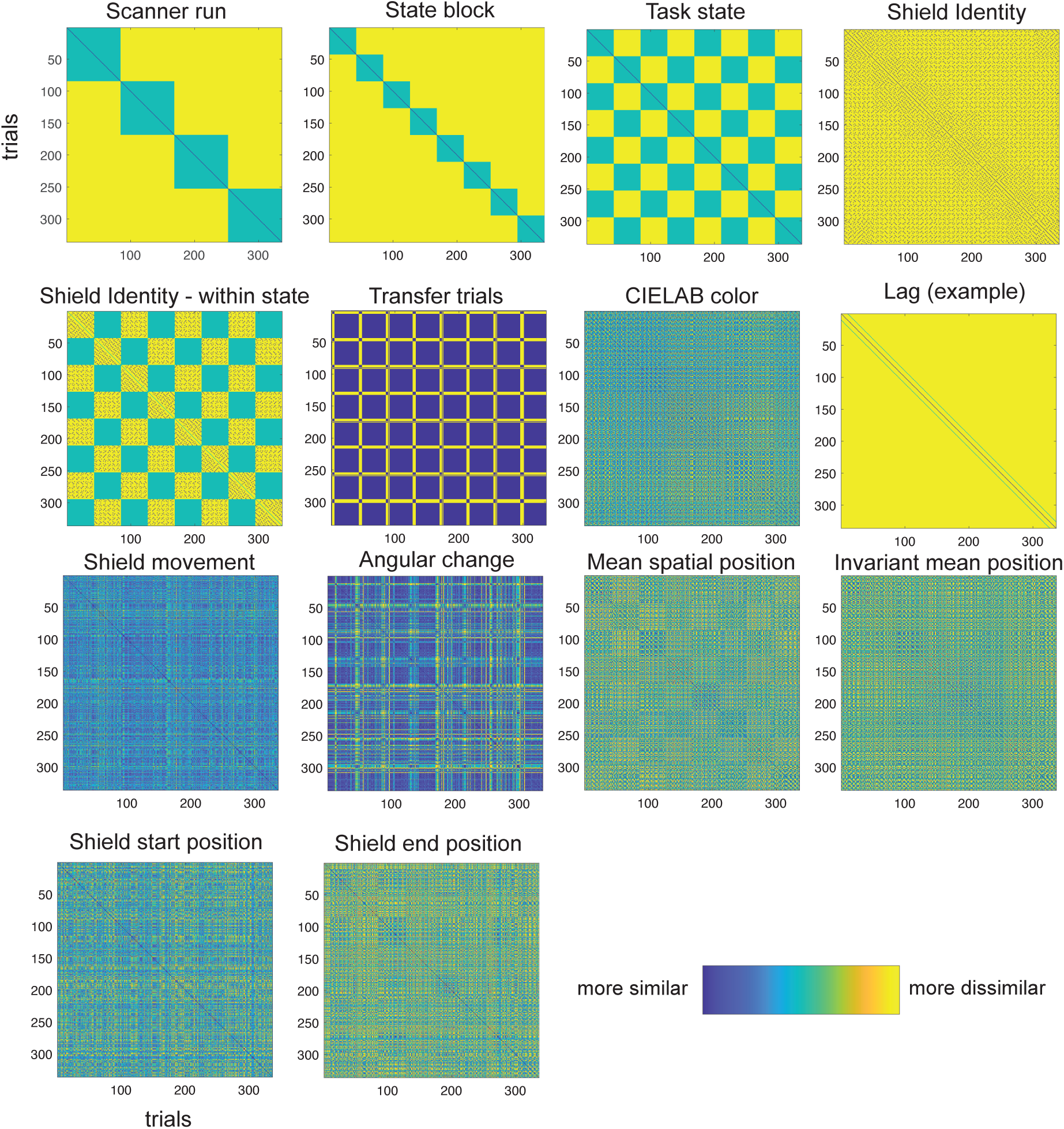
Hypothesis trial-wise representational dissimilarity matrices (RDMs) for representational similarity analysis from one randomly chosen example participant.

**Supplementary Figure 10.**
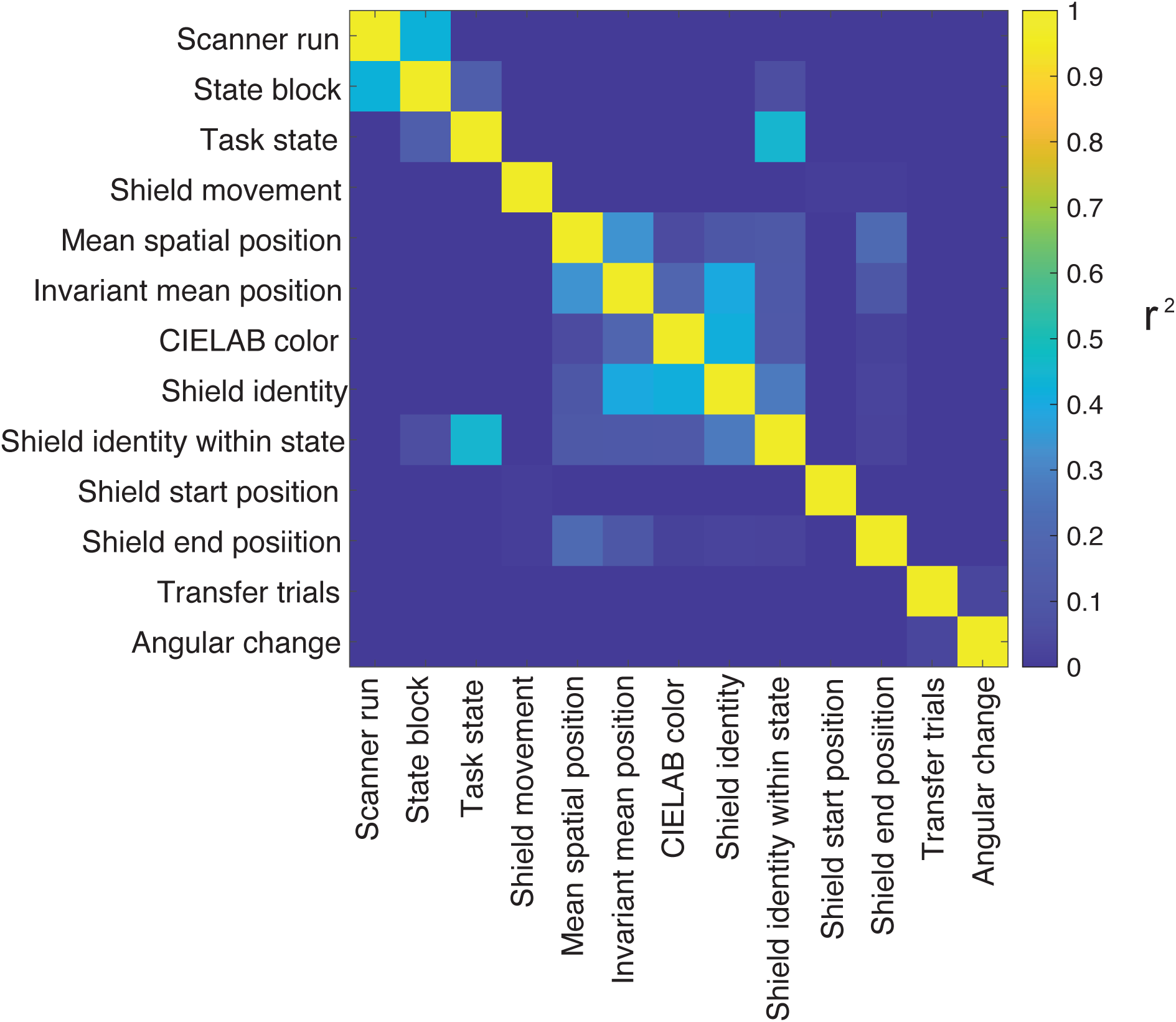
Variance shared (Pearson’s r^2^) between regressors included in representational similarity analysis, excepting lag regressors.

**Supplementary Figure 11.**
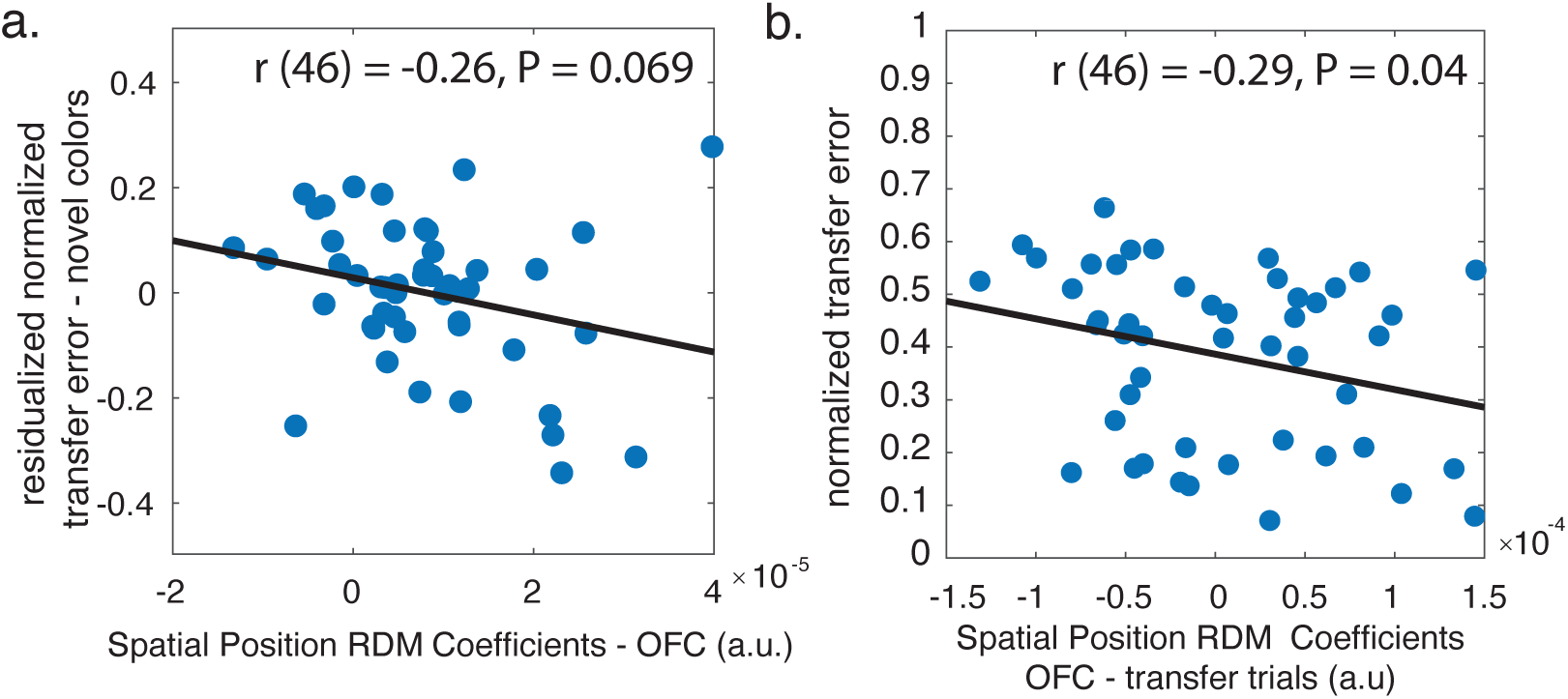
Additional tests of relationship between behavior and spatial position representation in bilateral orbitofrontal cortex (OFC). **a.** Scatterplot showing relationship between coefficients for mean spatial position representational dissimilarity matrix (RDM) in a ROI of OFC and normalized error on novel color transfer trials, after statistically removing the linear relationship between normalized error on these trials and normalized error on repeat trials for the original colors. **b.** Relationship of mean spatial position RDM coefficients in OFC for original color transfer trials plotted against normalized transfer error for the same.

## Notes

https://github.com/learning-memory-and-decision-lab/GridPlanetStudy

### Competing Interest Statement

The authors have declared no competing interest.

